# SARS-CoV-2 infection leads to acute infection with dynamic cellular and inflammatory flux in the lung that varies across nonhuman primate species

**DOI:** 10.1101/2020.06.05.136481

**Authors:** Dhiraj Kumar Singh, Shashank R. Ganatra, Bindu Singh, Journey Cole, Kendra J. Alfson, Elizabeth Clemmons, Michal Gazi, Olga Gonzalez, Ruby Escobedo, Tae-Hyung Lee, Ayan Chatterjee, Yenny Goez-Gazi, Riti Sharan, Rajesh Thippeshappa, Maya Gough, Cynthia Alvarez, Alyssa Blakley, Justin Ferdin, Carmen Bartley, Hilary Staples, Laura Parodi, Jessica Callery, Amanda Mannino, Benjamin Klaffke, Priscilla Escareno, Roy N. Platt, Vida Hodara, Julia Scordo, Adelekan Oyejide, Dharani K. Ajithdoss, Richard Copin, Alina Baum, Christos Kyratsous, Xavier Alvarez, Bruce Rosas, Mushtaq Ahmed, Anna Goodroe, John Dutton, Shannan Hall-Ursone, Patrice A. Frost, Andra K. Voges, Corinna N. Ross, Ken Sayers, Christopher Chen, Cory Hallam, Shabaana A. Khader, Makedonka Mitreva, Timothy J. C. Anderson, Luis Martinez-Sobrido, Jean L. Patterson, Joanne Turner, Jordi B. Torrelles, Edward J. Dick, Kathleen Brasky, Larry S. Schlesinger, Luis D. Giavedoni, Ricardo Carrion, Deepak Kaushal

**Affiliations:** Southwest National Primate Research Center; Texas Biomedical Research Institute, San Antonio, TX, 78227; Regeneron Pharmaceuticals, Inc., Tarrytown, NY 10591; Washington University in St Louis School of Medicine, St Louis, MO; Veterinary Imaging Consulting of South Texas, San Antonio, TX, 78258

**Keywords:** COVID-19, SARS-CoV-2, nonhuman primates, rhesus macaques, baboons, marmosets, animal models, BAL, CT

## Abstract

There are no known cures or vaccines for COVID-19, the defining pandemic of this era. Animal models are essential to fast track new interventions and nonhuman primate (NHP) models of other infectious diseases have proven extremely valuable. Here we compare SARS-CoV-2 infection in three species of experimentally infected NHPs (rhesus macaques, baboons, and marmosets). During the first 3 days, macaques developed clinical signatures of viral infection and systemic inflammation, coupled with early evidence of viral replication and mild-to-moderate interstitial and alveolar pneumonitis, as well as extra-pulmonary pathologies. Cone-beam CT scans showed evidence of moderate pneumonia, which progressed over 3 days. Longitudinal studies showed that while both young and old macaques developed early signs of COVID-19, both groups recovered within a two-week period. Recovery was characterized by low-levels of viral persistence in the lung, suggesting mechanisms by which individuals with compromised immune systems may be susceptible to prolonged and progressive COVID-19. The lung compartment contained a complex early inflammatory milieu with an influx of innate and adaptive immune cells, particularly interstitial macrophages, neutrophils and plasmacytoid dendritic cells, and a prominent Type I-interferon response. While macaques developed moderate disease, baboons exhibited prolonged shedding of virus and extensive pathology following infection; and marmosets demonstrated a milder form of infection. These results showcase in critical detail, the robust early cellular immune responses to SARS-CoV-2 infection, which are not sterilizing and likely impact development of antibody responses. Thus, various NHP genera recapitulate heterogeneous progression of COVID-19. Rhesus macaques and baboons develop different, quantifiable disease attributes making them immediately available essential models to test new vaccines and therapies.

## Main

A novel coronavirus, designated severe acute respiratory syndrome coronavirus 2 (SARS-CoV-2), emerged in Wuhan, China in 2019, and was proven to be the cause of an unspecified pneumonia. It has since spread globally, causing Coronavirus Disease 2019 (COVID-19) ^1^. The World Health Organization (WHO) declared COVID-19 a pandemic. It is clear that community spread of SARS-CoV-2 is occurring rapidly and the virus has very high infectivity and transmission rates, even compared to SARS-CoV-1, the causative agent of an outbreak 15 years earlier. It has been estimated that between up to 250,000 American lives may be lost due to COVID-19. The world over, these numbers could be 10-50 times worse. Clearly, COVID-19 is the most defining pandemic of this era, requiring significant biomedical research input, in order to most effectively fast track the development of new therapies and vaccines.

Human COVID-19 disease presents with a broad clinical spectrum ranging from asymptomatic to mild and severe cases. Patients with COVID pneumonia exhibit high-grade pyrexia, fatigue, dyspnea and dry cough accompanied by a rapidly progressing pneumonia, with bilateral opacities on x-ray and patchy, ground glass opacities on lung Computed Tomography (CT) scans. Individuals with immunocompromised conditions and comorbidities are at highest risk for worse outcomes of COVID-19.

Nonhuman primate (NHP) models of infectious diseases have proven useful for both investigating the pathogenesis of infection and testing therapeutic and vaccine candidates ^2^. During the SARS and MERS outbreaks, NHP models were developed with a moderate degree of success^3^. Early reports also indicate the utility of NHPs for SARS-CoV-2 infection, and for evaluating vaccine candidates^4, 5, 6, 7^. We hypothesized that the heterogeneity of human responses to SARS-CoV-2 infection can be recapitulated using multiple NHP species. Furthermore, we sought to gain a detailed characterization of the early cellular immune events following SARS-CoV-2 infection in the lung compartment, which has not yet been reported. Here, we compare SARS-CoV-2 infection in three species of NHPs (Specific Pathogen-free [SPF] Indian rhesus macaques, African-origin baboons, and New-World origin common marmosets). We assess age as a variable and focus our studies on high resolution imaging and the critical nature of the early cellular immune response in the lung which likely impacts disease outcome.

### Early events in SARS-CoV-2 infection in rhesus macaques

We first assessed the ability of SARS-CoV-2 to infect rhesus macaques during an acute 3-day infection study. Four Indian-origin mycobacteria- and SPF-naïve rhesus macaques (*Macaca mulatta*) (Table S1) were infected by multiple routes (ocular, intratracheal and intranasal) with sixth-passage virus at a target dose of 1.05×10^6^ PFU/per animal. All animals developed clinical signs of viral infection as evidenced by a doubling of serum C-Reactive Protein (CRP) levels relative to baseline, indicating systemic inflammation (Fig 1a); significantly decreased serum albumin (Fig 1b) and hemoglobin (Fig 1c) levels, indicating viral-induced anemia; and progressively increasing total serum CO_2_ levels (Fig S1a) indicative of pulmonary dysfunction. These observations were accompanied by a decrease in red blood cells (RBCs) (Fig S1b), reticulocytes (Fig S1c), white blood cells (WBCs) (Fig S1d), and platelet counts (Fig S1e); and a decrease in both the total number and percentage of neutrophils (Fig S1f, g), the latter suggesting that neutrophils are recruited to the lung compartment in response to SARS-CoV-2 infection as first responders. In contrast, systemic influx of monocytes was observed, indicating viral infection-induced myelopoiesis (Fig S1h). Monocytes are crucial for successful antiviral responses via recognition of pathogen-associated molecular patterns, thereby initiating a signaling cascade that invokes an interferon response to control infections. No significant pyrexia or weight loss was observed in this acute study. Overall, our results suggest that rhesus macaques develop several clinical signs of viral infection following experimental exposure to SARS-CoV-2.

**Figure 1.**
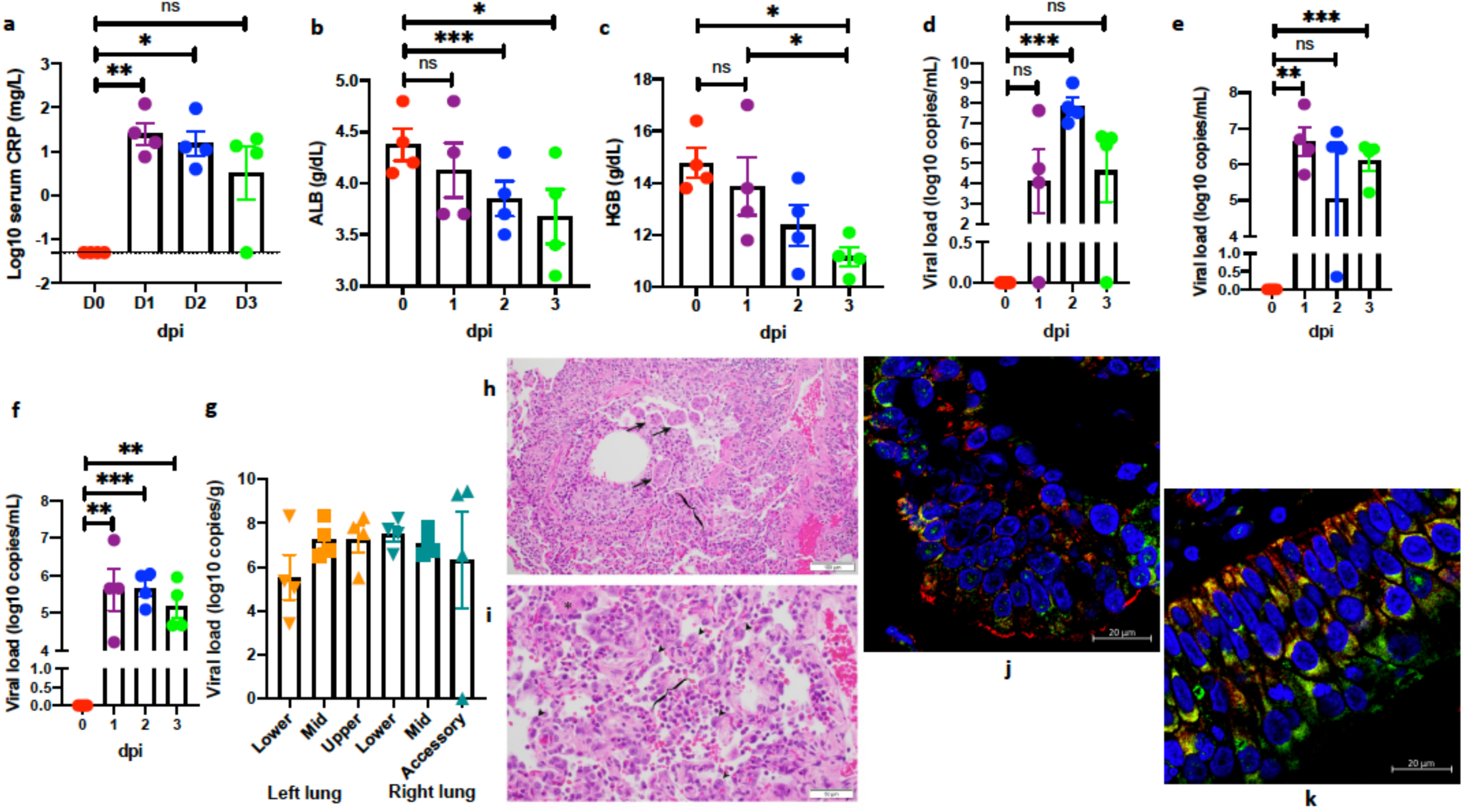
**Clinical correlates of SARS-CoV-2 infection in rhesus macaques over 0-3 dpi**. Changes in serum CRP (mg/L) (a), albumin (ALB) (g/dL) (b), hemoglobin (HGB) content (g/dL) (c), longitudinally in peripheral blood. Viral RNA (log_10_ copies/mL were measured by RT-PCR in BAL fluid (d), nasopharyngeal (e), and buccopharyngeal (f) swabs longitudinally (red – 0 dpi; purple – 1 dpi; blue – 2 dpi; green – 3 dpi). Viral RNA was also measured in lung tissue homogenates at endpoint (3 dpi) and data is expressed as log_10_ copies/gram of the lung tissue for random samples from three lobes in left (orange) and right (teal) lungs (g Hematoxylin and eosin (H&E) staining was performed on formalin-fixed paraffin-embedded (FFPE) lung sections from infected animals for pathological analysis Histopathologic analysis revealed bronchitis characterized by infiltrates of macrophages, lymphocytes, neutrophils, and eosinophils that expanded the wall (bracket), and along with syncytial cells (arrows) filled the bronchiole lumen and adjacent alveolar spaces. (h); Suppurative interstitial pneumonia with Type II pneumocyte hyperplasia (arrowheads) and alveolar space filled with neutrophils, macrophages and fibrin (*). Bracket denotes alveolar space. (i). Multilabel confocal immunofluorescence microscopy of lungs (j) and nasal epithelium (k) at 63x with Nucleocapsid (N) specific antibody (green) DAPI (blue), and ACE2 (red). (a-f) Data is represented as mean± SEM (n=4).). (c-g) Undetectable results are represented as 1 copy. One way Repeated-measures ANOVA with Geisser-Greenhouse correction for sphericity and Tukey’s post hoc correction for multiple-testing (GraphPad Prism 8) was applied. * P<0.005, ** P<0.005, *** P<0.0005.

Viral RNA was detected in BAL, and from nasal or nasopharyngeal (NS) and buccopharyngeal (BS) swabs at 1-3 days post-infection (dpi), but not at pre-infection time points (Fig 1d-f). Viral RNA was also detected in saliva and from rectal swabs (RS) in a small subset of animals (Fig S1i-j). Unlike other samples, viral RNA was only detected in RS at later time points (i.e., after 1 dpi). At necropsy (3 dpi), we performed random sampling from every lung lobe and SARS-CoV-2 RNA could be detected in 23/24 total lung sections analyzed. An average of 4-6 log copies/100 mg of lung tissue could be detected from every lobe (Fig 1g). The ∼4-log increase in viral RNA from 1 to 2 dpi in the BAL (Fig 1d) provided clear evidence of early active replication of SARS-CoV-2 in rhesus macaques.

Examination at necropsy (3 dpi) revealed findings of interstitial and alveolar pneumonia (Fig 1h, i). While gross appearance of the lungs of most infected animals was unremarkable (Fig S2a), multifocal to coalescing red discoloration of the left lung lobes in one macaque was observed (Fig S2b). Table S2 summarizes the histopathologic findings in descending order of occurrence by anatomic location. The lung was the most affected organ ((Fig 1h, i, Table S2, Fig S2). Multifocal, mild to moderate interstitial pneumonia characterized by infiltrates of neutrophils, macrophages, lymphocytes, and eosinophils was present in all four animals (Fig 1 i, Fig S2d, e, g, h), and was accompanied by variable fibrosis (4/4, Fig S2e), fibrin deposition (3/4, Fig S2c), vasculitis (3/4, Fig S2f), edema (2/4, Fig S2h), necrosis (Fig S2g), and areas of consolidation (2/4, Fig S2c). All four macaques exhibited the following: 1) Syncytial cells in the epithelial lining and/or alveolar lumen (Fig S2e, g, k); 2) Bronchitis characterized by infiltrates of eosinophils within the bronchial wall and epithelium (Fig 1h, Fig S2i, j, k); Bronchus-associated lymphoid tissue (BALT) hyperplasia (Fig S2i); and 4) Minimal to moderate lymphoplasmacytic and eosinophilic tracheitis and rhinitis.

The presence of SARS-CoV-2 in tissue sections collected at necropsy (3 dpi) was determined by multi-label confocal immunofluorescence using antibodies specific for Nucleocapsid (N) (Fig 1j, k, Fig S3) and Spike(S) proteins (Fig S4) and their respective isotype controls (Fig S3, S4). Fluorescence immuno-histochemical analysis revealed the presence of SARS CoV-2 proteins in lungs (Fig 1j, Fig S3a, g, Fig S4a, d, g, j), nasal epithelium (Fig 1k, Fig S3b, h, Fig S4b, e, h, k) and tonsils (Fig S3c, i, Fig S4c, f, I, l). In all tissues, including lungs (Fig 1j, Fig S3 a, g), nasal epithelium (Fig 1k, Fig S3 b, h) and tonsils (Fig S3 c, i), N antigen signal was present in cells expressing ACE2, which has been shown to be a receptor for SARS-CoV-2, or in cells adjoining those expressing ACE2. No signal was detected in N isotype control staining in lungs, nasal epithelium or tonsils (Fig S3d-f), and no signal for viral antigen was detected in naïve tissues (Fig S3 m, n). It appeared that the expression levels of ACE2 protein were much lower in lung tissues derived from naïve animals compared to those from macaques exposed to SARS-CoV-2 (Fig S3 m, n). The majority of the S signal was detected in the epithelial layer with discrete distribution throughout the lung tissue (Fig S4a, d). In the nasal cavity, the virus was observed in cells of the epithelial linings (Fig S4b, h) but in tonsils, the virus appeared distributed throughout the tissue (Fig 3c, i). Together, these results show that SARS-CoV-2 exposure induces a respiratory tract infection in rhesus macaques. Viral replication is supported in the upper and lower lung compartments during the first three days of infection and viral antigens are detected at high levels in the lungs.

To complement the lung histopathology in rhesus macaques, radiographs were performed at baseline and each day post-infection. All four infected macaques showed progressive increase in CXR abnormality scores, consistent with an infectious disease (Fig 2a, Fig S5a). The 2 and 3 dpi CXR scores were significantly elevated relative to baseline (Fig 2a), despite evidence of partial resolution of specific lesions at 2 or 3 dpi versus 1 dpi (Fig S5a). There were mild-to-severe multifocal interstitial-to-alveolar patterns with soft tissue opacities (seen as ground glass opacities described in the CT scans below) in various lobes or diffusely in some animals, with more severe abnormalities in the lower lung lobes, and with the most severe findings at 3 dpi (Fig S5a). Pleural effusions were also observed.

**Figure 2.**
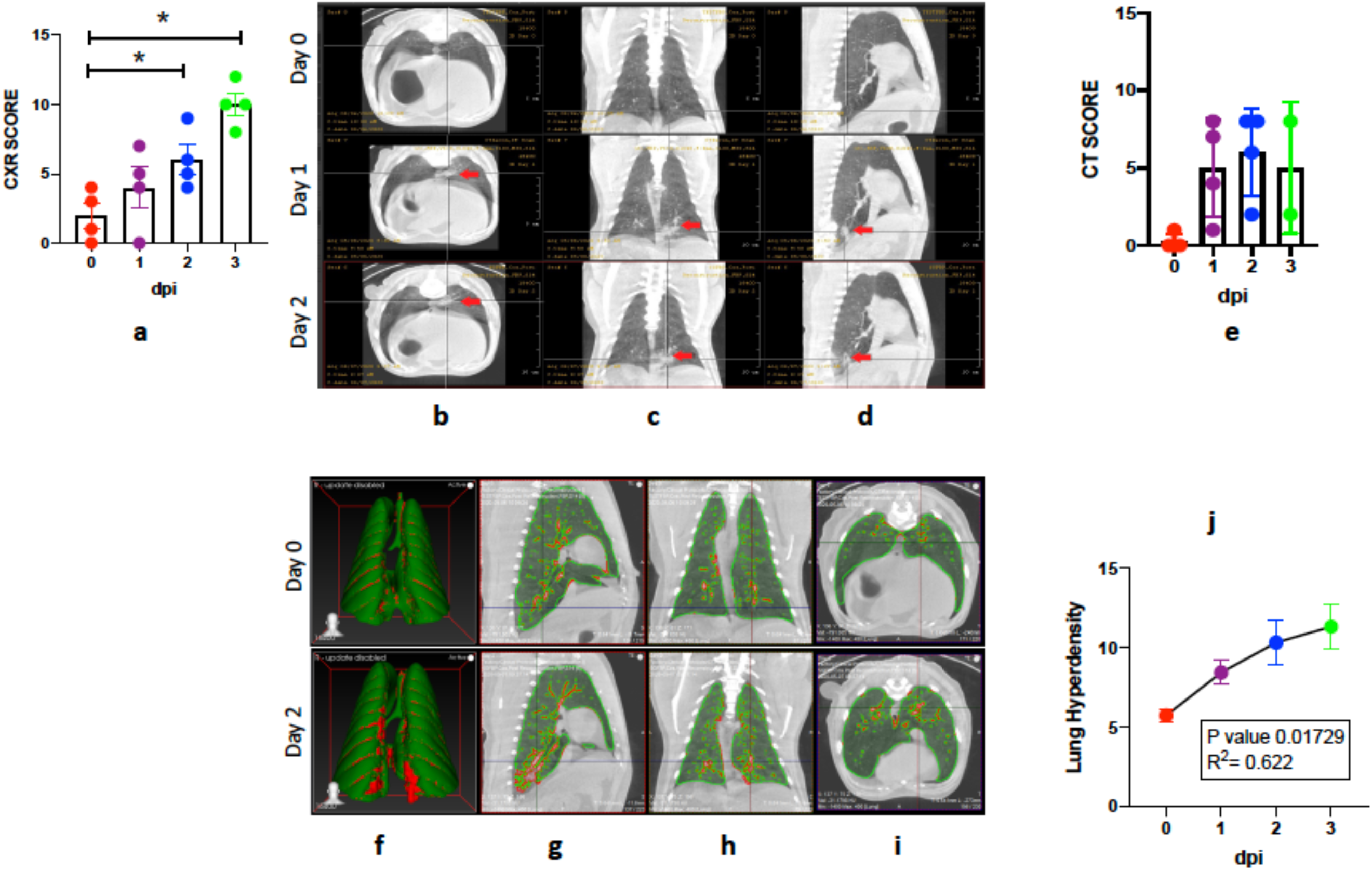
**Radiologic evaluation of the lung compartment following SARS-CoV-2 infection in rhesus macaques over 0-3 dpi including by hyperdensity analyses**. CXR (a) and CT (e) scores generated by a veterinary radiologist blinded to the experimental group (red – 0 dpi; purple – 1 dpi; blue – 2 dpi; green – 3 dpi). (a) Data is represented as mean± SEM (n=4). One way Repeated-measures ANOVA with Geisser-Greenhouse correction for sphericity and Tukey’s post hoc correction for multiple-testing (GraphPad Prism 8) was applied. * P<0.05. Representative CT scan images performed on Day 0-2 dpi show (b) transverse, (c) vertical, (d) longitudinal view of left caudal lobe ground glass opacity on 1 dpi (middle), 2 dpi ^28^ and baseline at 0 dpi (upper inset). CT scans (b-d) revealed evidence of pneumonia and lung abnormalities in the infected animals relative to controls which resolved between 1 to 2 dpi (red arrow). 3D reconstruction (f) of ROI volume representing the location of lesion. (Fig 2g-i) represent image for quantification of lung lesion with green area representing normal intensity lung voxels (−850 HU to −500 HU), while red areas represent hyperdense voxels (−490 HU to 500 HU). Percent change in lung hyperdensity in SARS-CoV2 infected animals over Day 1-3 dpi compared to the baseline(j). (red – 0 dpi; purple – 1 dpi; blue – 2 dpi; green – 3 dpi). (e, j) Data represented as (mean ± SEM) (n=4 for 0-2 dpi, n=2 for 3dpi). Ordinary one-way ANOVA with Dunnett’s post hoc test was applied.

Lung CT scans prior to infection showed a normal thorax cavity with the exception of atelectasis (Fig 2b-d, top panel). Within 1 dpi, CT scans showed increased multifocal pulmonary infiltrates with ground glass opacities in various lung lobes, linear opacities in the lung parenchyma, nodular opacities in some lung lobes, and increased soft tissue attenuation extending primarily adjacent to the vasculature (Fig 2c, Fig S5b-e). In some animals, multifocal alveolar pulmonary patterns and interstitial opacities were observed in lobe subsections, with soft tissue attenuation and focal border effacement with the pulmonary vasculature. Features intensified at 2-3 dpi, primarily in the lung periphery, but also adjacent to the primary bronchus and the vasculature (Fig 2d, Fig S5b). In other animals, progressive alveolar or interstitial pulmonary patterns were observed at 2 dpi (Fig 2c). While ground glass opacities in some lobes intensified at 2 dpi relative to 1 dpi, others resolved (Fig S5c, d). In one animal, the individual nodular pattern at 1 dpi evolved to a multifocal soft tissue nodular pattern in multiple lobes with associated diffuse ground glass opacities (Fig S5d). At 3 dpi, persistent, patchy, fairly diffuse ground glass pulmonary opacities existed in many lung lobes with multifocal nodular tendency (Fig S5e). Overall, CT abnormality scores continuously increased at over the 3 days relative to baseline (Fig 2e). Percent change in the hyperdensity volume was calculated using CT scans to quantify pathological changes over the course of disease ^8^. We observed a significant increase in lung hyperdense areas between 1-3 dpi compared to the baseline scans (Fig 2f-i). Measurement of volume involved in hyperdensity showed a significant, progressive increase over time (Fig 2j). Pneumonia was evident in all infected animals relative to their baseline (Fig 2j), suggesting that while some lesions formed and resolved within the three-day infection protocol, others persisted or progressed. Together, CXR and CT scans revealed moderate multi-lobe pneumonia in infected animals, confirming the histopathology results (Fig 1h, i, Fig S2) in the very early phase of SARS-CoV-2 infection in rhesus macaques.

We measured the levels of pro-inflammatory, Type I cytokines in the BAL fluid (Fig 3) and plasma (Fig S6a-l) of acutely infected rhesus macaques. Levels of IL-6 (Fig 3a), IFN-*α* (Fig 3b), IFN-*γ* (Fig 3c), IL-8 (Fig 3d), perforin (Fig 3e), IP-10 (Fig 3f), MIP1-*α* (Fig 3g) and MIP1-*β* (Fig 3h) were all significantly elevated in the BAL fluid. The levels of IL-12p40 (Fig 3i), IL-18 (Fig 3j), TNF (Fig 3k) and IL-1Ra (Fig 3l) increased over time. Of particular interest was the elevation of Type I IFN-*α* (Fig 3b), which has critical anti-viral activity including against SARS-CoV-2 ^9^. Expression of a downstream Type-I interferon-regulated gene IP-10 (CXCL-10), which promotes the recruitment of CXCR3^+^ Th1 thymocytes, was also induced (Fig 3f). Therefore, we observed that rhesus macaques mount an early anti-viral response to SARS-CoV-2 infection. Type I IFNs and IL-6 (both significantly expressed) are key components of a “cytokine-storm” which promote acute respiratory distress syndrome (ARDS) associated with both SARS-CoV-1 and -2, when induced uncontrollably^10^. IFN-*α* and IP-10 were also significantly elevated in plasma samples at 2 and 3 dpi (Fig S6).

**Figure 3.**
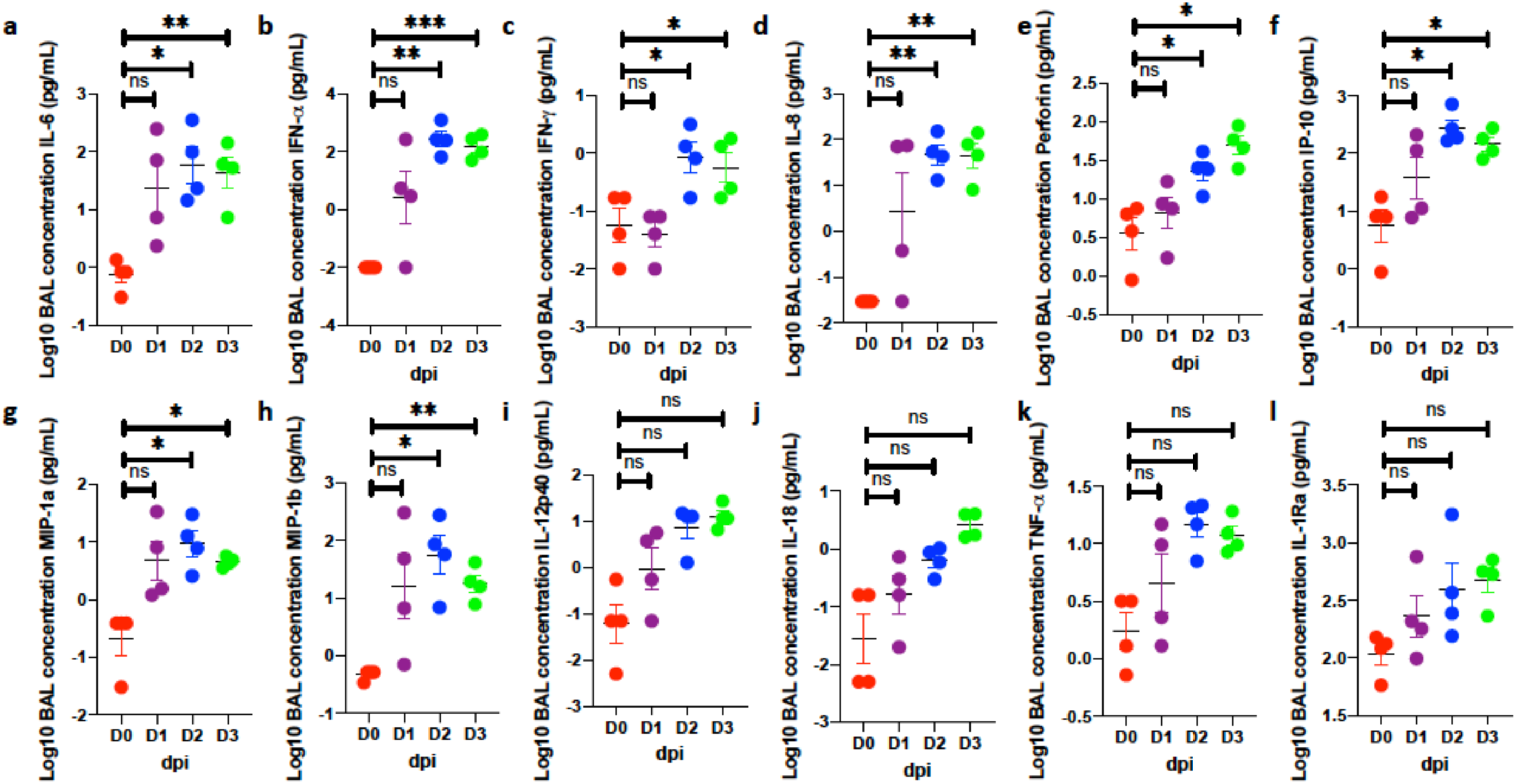
**SARS-CoV-2 induced alveolar inflammation**. Simultaneous analysis of multiple cytokines by Luminex technology in the BAL fluid of rhesus macaques over 0-3 dpi. Levels of IL-6 (a), IFN-a (b), IFN-g (c), IL-8 (d), perforin (e), IP-10 (f), MIP1a (g), MIP1b (h), IL-12p40 (i), IL-18 (j), TNF (k) and IL-1Ra (l) are expressed in Log10 concentration in picogram per mL of BAL fluid. (red – 0 dpi; purple – 1 dpi; blue – 2 dpi; green – 3 dpi). Data is represented as mean± SEM (n=4). One way Repeated-measures ANOVA with Geisser-Greenhouse correction for sphericity and Tukey’s post hoc correction for multiple-testing (GraphPad Prism 8) was applied. * P<0.005, ** P<0.005, *** P<0.0005.

Thus, clinical, imaging, pathology and cytokine analyses provide evidence for an acute infection in macaques following exposure to SARS-CoV-2, which leads to a moderate pneumonia and pathology, with early activation of anti-viral responses. To study progression of infection, and assess the effect of age on SARS-CoV-2 infection, we infected six young and six old Indian-origin rhesus macaques as described above and longitudinally followed the outcome over 14-17 days (Table S1). We included four macaques as procedural controls, which were sham-infected and underwent all procedures (with the exception of necropsy) to control for the impact of multiple procedures over the course of the study (Table S1). We also infected six baboons and an equal number of marmosets with SARS-CoV-2 in order to compare the progression of COVID-19 in different NHP models.

### Long-term study of SARS-CoV-2 infection in rhesus macaques, baboons and marmosets demonstrates heterogeneity in progression to COVID-19

Results of the longitudinal study showed that the acute signs of SARS-CoV-2 infection and mild-to-moderate COVID-19 disease in rhesus macaques markedly improved over time (Fig S7). In general, no major differences were observed as a consequence of age, and subsequent data from young and old animals are combined (N=12), unless specified. A small subset (3/12) of animals exhibited elevated serum CRP past 3 dpi (Fig S7a), although metabolic signs of dysfunction likely induced by infection (e.g. tCO2 elevation) continued for the duration of the study (Fig S7b). No alterations were observed in the levels of serum albumin or hemoglobin during this timeframe (not shown). There was a significant decline in RBCs (Fig S7c) at 3 and 6 dpi which normalized or reverted by 9 dpi. The percentage of neutrophils in the peripheral blood remained unchanged between 3-14 dpi (not shown). The significant decline in blood platelets and increase in the percentage of monocytes observed at 3 dpi, were short-lived (Fig S7d, e). Despite these modest changes, the majority of animals in both age groups exhibited weight loss throughout the study duration (Fig S7f), although pyrexia was not observed (not shown).

Viral RNA was detected in BAL of 10/12 macaques at 3 dpi, but declined thereafter (Fig 4a). Detection of viral RNA was equivalent between young (5/6) and old (5/6) macaques (Fig S8a). Very low viral RNA copy numbers were detected in BAL at 9 dpi with only one young macaque testing positive, and none by 12 dpi (Fig 4a). Viral RNA appeared to persist for much longer in NS than BAL, including at study endpoint (Fig 4b). Viral RNA was detected from NS in 6/12 macaques at 3 dpi and on average young macaques harbored more virus in their nasal cavity at 3 dpi relative to old animals but the differences were not significant (Fig S8b). SARS-CoV-2 RNA was detected in 10/12 macaques (6 young, 4 old, respectively) at 9 dpi and 6/12 macaques at the end of the study period (Fig S8b). These results suggest that the virus persists for at least two weeks in the respiratory compartment of immunocompetent macaques that clinically recovered from COVID-19. Viral RNA was detected from BS in 4/12 animals at 3 and 6 dpi, but not at later time points (Fig S8c, d). No significant difference was detected between age groups. Viral RNA was also detected from RS in 2/12 animals at 3, 6 and 9 dpi (Fig S8e, f). Significantly lower levels of viral RNA (2.5 logs) were detectable at the end of the study (14-17 days) when compared to viral RNA detected at the end of the 3-day protocol (Fig 4c, Fig S9a). Viral RNA was detected in the lungs of two-thirds (8/12) of all macaques and no effect of age was apparent. No viral RNA was detected in any serum samples (Fig S9b) or in randomly selected urine samples (Fig S9c). The presence of viral RNA in the lungs of macaques after two weeks following recovery from acute COVID-19 indicates that while macaques control SARS-CoV-2 infection, immune responses are not sterilizing.

**Figure 4.**
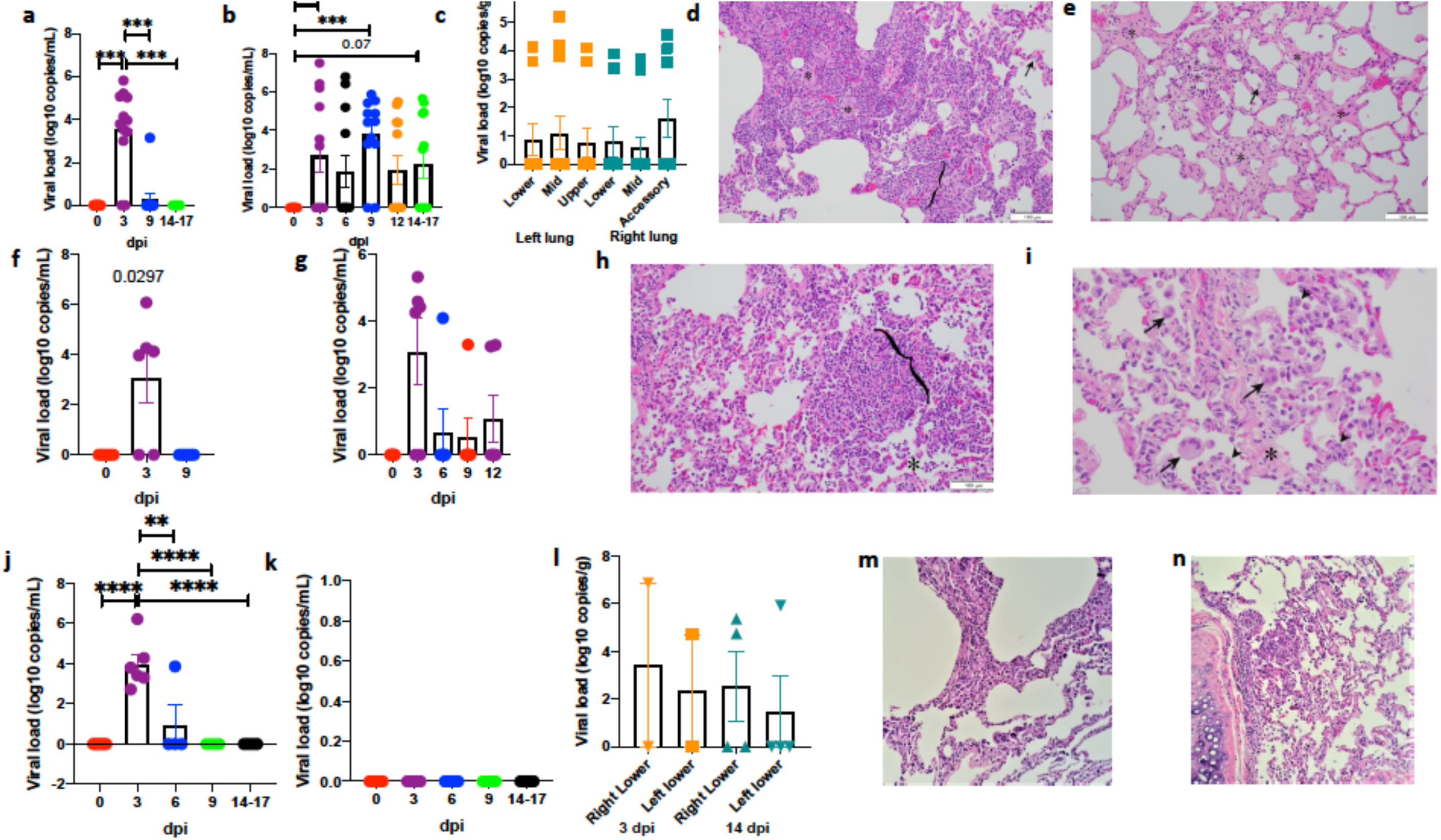
**Longitudinal clinical and histopathological correlates of SARS-CoV-2 infection in rhesus macaques, baboons and marmosets over two weeks**. Viral RNA (log_10_ copies/mL were measured by RT-PCR in BAL fluid (a) and nasopharyngeal (b) swabs of SARS-CoV-2 infected rhesus macaques longitudinally (red – 0 dpi; purple – 3 dpi; black – 6 dpi: blue – 9 dpi; orange – 12 dpi: green – 14-17 dpi). (n=12) One way Repeated-measures ANOVA with Geisser-Greenhouse correction for sphericity and Tukey’s post hoc correction for multiple-testing (GraphPad Prism 8) was applied. * P<0.005, *** P<0.0005. Viral RNA was also measured in lung tissue homogenates of infected rhesus macaques at endpoint (14-17 dpi) and data is expressed as log_10_ copies/gram of the lung tissue for random samples from three lobes in left (orange) and right (teal) lungs (c). Histopathologic analysis revealed regionally extensive interstitial lymphocytes, plasma cells, lesser macrophages and eosinophils expanding the alveolar septa (bracket) and alveolar spaces filled with macrophages (*). Normal alveolar wall is highlighted (arrow) for comparison (d). Alveolar spaces with extensive interstitial alveolar wall thickening by deposits of collagen (*) and scattered alveolar macrophages (arrow) (e). Viral RNA (log_10_ copies/mL were measured by RT-PCR in BAL fluid (f) and Nasopharyngeal (g) swab from SARS-CoV-2 infected baboons. (n=6) One way Repeated-measures ANOVA with Geisser-Greenhouse correction for sphericity and Tukey’s post hoc correction for multiple-testing (GraphPad Prism 8) was applied. Histopathologic analysis revealed regionally extensive interstitial lymphocytes, plasma cells, lesser macrophages and eosinophils expanding the alveolar septa (bracket) and alveolar spaces filled with macrophages (*), (h). Alveolar wall thickening by interstitial deposits of collagen (*), alveoli lined by occasional type II pneumocytes (arrowhead) and alveolar spaces containing syncytial cells (arrow) and alveolar macrophages (i). Viral RNA (log_10_ copies/mL were measured by RT-PCR in marmoset nasal wash (j) and oral (k) swabs longitudinally (red – 0 dpi; purple – 3 dpi; blue – 6 dpi; green – 9 dpi; black – 14-17 dpi). n=6 for 0-3 dpi and n=4 for 6-14 dpi). Histopathologic analysis revealed milder form of interstitial lymphocytes, and macrophages recruited to the alveolar space (m, n). Ordinary one-way ANOVA with Dunnett’s post hoc test was applied. Viral RNA was also measured in lung homogenates at endpoint (3 dpi & 14 dpi) and data is expressed as log_10_ copies/gram of the lung for random samples from left and right lobes at 3 dpi (orange) and 14 dpi (teal) (g). Data is represented as mean± SEM. ** P<0.005, **** P<0.00005.

Gross examination of the lungs of most infected animals at necropsy (14 to 17 dpi) was unremarkable (Fig S10a); however, red discoloration of the dorsal aspect of the lung lobes was seen in four young and two aged animals (Fig S10b). Table S3 summarizes the histopathologic findings in descending order of occurrence by anatomic location. The lungs were the most affected organ (Fig 4d, e, Table S3). Multifocal minimal to mild interstitial mononuclear inflammation was seen in 11/12 animals (Fig 4n, 0, Fig S10c), generally composed of macrophages and lymphocytes that expanded the alveolar septa (Fig 4n, Fig S10d, e, f, g), with variable neutrophil infiltrates (5/12, Fig S10e), fibrosis (5/12, Fig 4o, Fig S10f, g) or vasculitis (3/12, Fig S10i). Alveolar epithelium often contained areas of type II pneumocyte hyperplasia (4/12, Fig S10e) and bronchiolization (2/12, Fig S10h). Alveolar lumina contained increased alveolar histiocytosis (9/12, Fig 4o, Fig S10d, e) occasionally admixed with neutrophils (5/12, Fig S10e). Syncytial cells (Fig S10e, f) were observed most frequently in the alveolar lumen in all 12 animals. Bronchitis was observed in 4/12, characterized by infiltrates of eosinophils within the bronchial wall and epithelium (Fig S10j). Prominent perivascular lymphocytes (7/12, Fig S10k) and BALT hyperplasia (5/12, Fig S10i) were frequently observed. The majority of the animals (11/12) exhibited minimal to moderate lymphoplasmacytic and eosinophilic tracheitis.

Early detection of viral RNA in BAL was comparable between macaques and baboons (Fig 4a, f) and NS (Fig 4b, g). A third of the baboons had detectable viral RNA in NS at 12 dpi (Fig 4g), and a similar number of animals remained positive at 9 dpi in the BS (Fig S11a). The number of baboons from which viral RNA could be detected in RS increased over time from 1/6 at 3 dpi to 3/6 at 6 dpi, 4/6 at 9 dpi and 3/6 at 12 dpi, underscoring long-term viral persistence of SARS-CoV-2 in baboons relative to rhesus macaques (Fig S11b). Postmortem gross examination at 14 to 17 dpi identified red discoloration of the lung lobes in all six baboons (Fig S11c, d). Table S4 summarizes the histopathologic findings in descending order of occurrence by anatomic location. Like macaques, the lungs were the most affected organ in the baboons (Fig 4h, i, Table S6, Fig S11). Multifocal minimal to moderate interstitial mononuclear inflammation was seen in 6/6 animals (Fig 4h, Fig S11f, g, h, i), generally composed of macrophages and lymphocytes that expanded the alveolar septa, with variable neutrophil infiltrates (3/6, Fig S11f, g, j, k) or fibrosis (2/6, Fig S11j, k). Alveolar epithelium often contained areas of type II pneumocyte hyperplasia (4/6, Fig 4i, Fig S11i) and bronchiolization (1/6, Fig S11l). Alveolar lumina contained increased alveolar histiocytosis (6/6, Fig 4h) occasionally admixed with neutrophils (3/6) (Fig S11f, g, h, i). Syncytial cells were observed most frequently in the alveolar lumen in all 6 animals (Fig 4h, Fig S11m). Bronchitis was observed in 6/6, characterized by infiltrates of eosinophils within the bronchial wall and epithelium (Fig S11n). BALT hyperplasia (5/6, Fig S11k) was frequently observed. The majority of the animals exhibited minimal to moderate lymphoplasmacytic and eosinophilic tracheitis (5/6) and rhinitis (4/6).

SARS-CoV-2 infection was milder in marmosets. Less than 4 logs of viral RNA could be detected in NS from infected marmosets, peaking at 3 dpi, and 1/6 animals was also positive at 6 dpi. No viral RNA was detected at later time points (Fig 4j). No viral RNA was detected in BS (Fig 4k). A subset of six marmosets was euthanized at 3 dpi (n=2), while others were necropsied at 14 dpi. Approximately 2 logs of viral RNA could be detected in the lungs of marmosets at both time points. Evidence of SARS-CoV-2 infection-induced pathology, including interstitial and alveolar pneumonitis was observed in marmoset lungs as well (Fig 4m, n), although not as prevalent as in macaques or baboons. Thus, our results show that three genera of NHPs develop different degrees of COVID-19 following SARS-CoV-2 infection when evaluated side by side, with baboons exhibiting moderate to severe pathology, macaques exhibiting moderate pathology and marmosets exhibiting mild pathology. Viral RNA levels in BAL, NS and lungs are consistent with the levels of pathology. While other results also suggest that marmosets are unaffected by SARS-CoV-2 infection ^4^ (https://www.biorxiv.org/content/10.1101/2020.03.21.001628v1), we show that these NHPs do develop non-negligible, mild COVID-19-related pathology and some degree of viral persistence.

We performed detailed imaging of macaques in the longitudinal study. Similar to the acute study, imaging revealed the development of viral pneumonia. All macaques infected with SARS-CoV-2 exhibited low baseline CXR scores (Fig 5a, Table S5) with no difference due to age (Fig 5b). Several infected macaques showed changes consistent with pneumonia (Table S5) with peak severity seen between 3-6 dpi, followed by a decline by study end (Fig 5a, b, Table S5). Examples of the development of extensive pneumonia by CXR can be seen in macaques at 6 dpi, relative to baseline with subsequent resolution (Fig 5 c-e). Several animals exhibited multi-lobe alveolar infiltrates and/or interstitial opacities at 6 dpi. In other animals, there were progressive, moderate to severe interstitial and alveolar infiltrates at 6 dpi, which resolved by day 14. Conversely, the radiographs of all procedure control animals (which underwent repeated BAL procedures) exhibited normal a thorax cavity with minimal to no findings.

**Figure 5.**
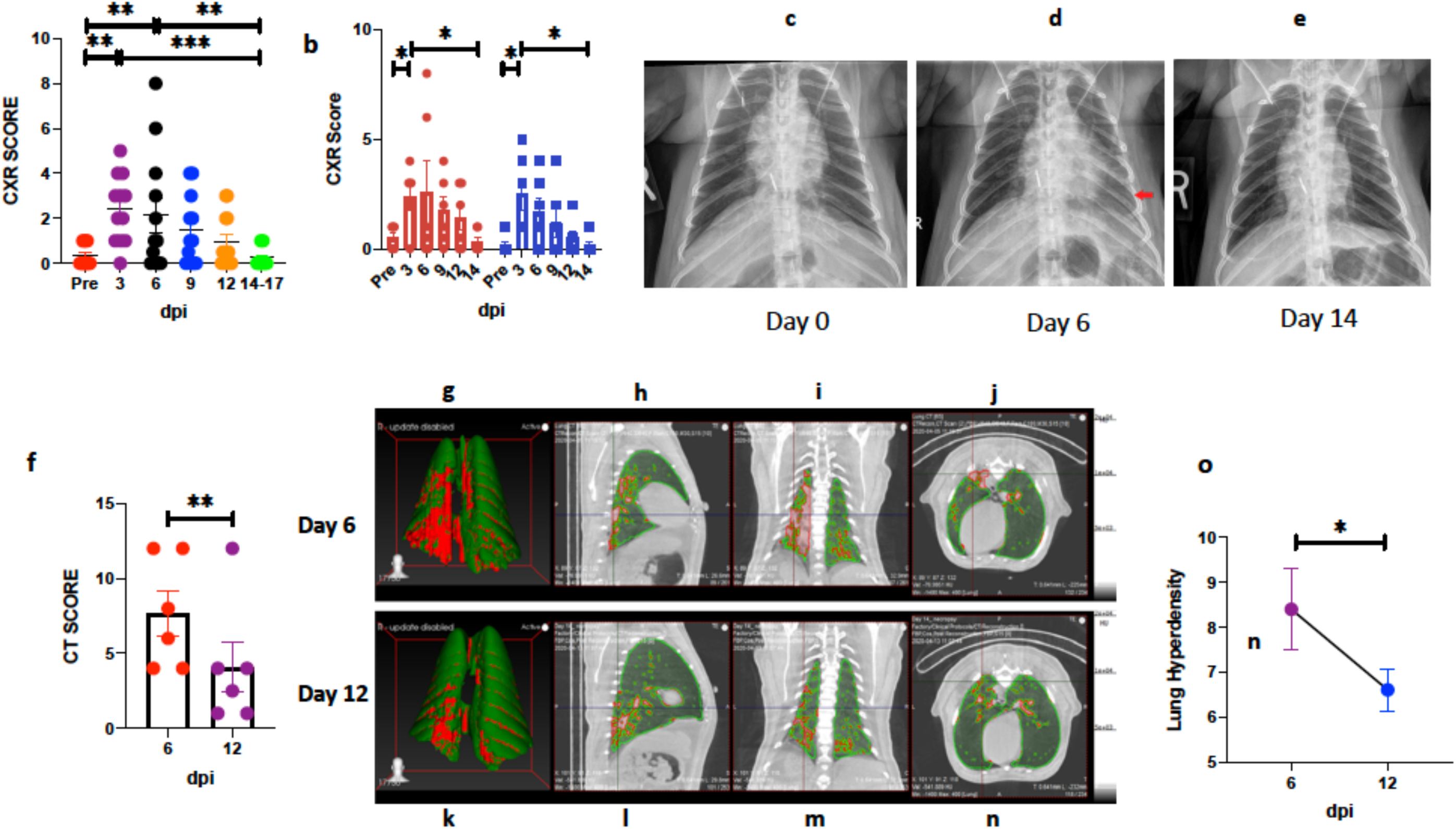
**CXR** (a) scores generated by a veterinary radiologist blinded to the experimental group (n=12) and (b) CXR scores split in old and young macaques (n=6). CXR radiographs showing minimal right caudal interstitial pattern at 0 dpi (c), Alveolar pattern associated with the caudal sub segment of the left cranial lung lobe and left caudal lung lobe with patchy right caudal interstitial opacity at 6 dpi (d) and Minimal left caudal interstitial pattern at 14dpi (e). CT (f) scores generated by a blinded veterinary radiologist (n=6). 3D reconstruction (g,k) of ROI volume representing the location of lesion. (h-j, l-n) represent image for quantification of lung lesion with green area representing normal intensity lung voxels (−850 HU to −500 HU), while red areas represent hyperdense voxels (−490 HU to 500 HU). Percent change in lung hyperdensity in SARS-CoV2 infected animals over 6 dpi compared to 12 dpi (o) (n=6). Data is represented as mean± SEM. (a) One way & (b) two way Repeated-measures ANOVA with Geisser-Greenhouse correction for sphericity and Tukey’s post hoc correction for multiple-testing and (f,o) Paired T test (GraphPad Prism 8) was applied. * P<0.005, ** P<0.005, *** P<0.0005.

High resolution CT imaging of the lungs was performed prior to and following SARS-CoV-2 infection on six young and six old macaques. Pneumonia was present in all animals, post-infection, but to a significantly higher degree in old macaques relative to young (Fig 5f, Fig S12a-f, Table S6). At 6 dpi, severe patchy alveolar patterns were observed in some lobes, while other lobes had milder, interstitial patterns, with moderate to severe ground glass opacities primarily in the lungs of old macaques (Fig S12a-f). In all animals, resolution of many ground glass opacities and nodular as well as multifocal lesions was observed at 12 dpi (Fig 5f, Fig S12a, b, d-f). At 12 dpi, all but one of the older macaques exhibited a normal or nearly normal thorax cavity, the latter with minimal ground glass opacities in all lung lobes studied at this time. Findings in one older macaque was considerably improved but retained patchy round glass opacities in all lobes and alveolar patterns in some lobes at 12 dpi (Fig S12c). This animal had the highest overall score by CT (Fig 5f) and CXRs (Fig 5 a-b). These results suggest that pneumonia in some older macaques may persist longer than in younger animals. Similar to the acute study, hyperdensity analysis revealed a significant, progressive increase in the volume of lung involved in pneumonia at 6 dpi, which normalized by 12 dpi (Fig 5g-o).

### SARS-CoV-2 infection in macaques results in a dynamic myeloid cell response in the lungs of rhesus macaques

Cellular composition in BAL samples and peripheral blood^11, 12^ at necropsy showed markedly altered immune cell responses in the lung compartment following infection of macaques. In healthy lungs, BAL is predominantly comprised of alveolar macrophages (AMs)^13^ but respiratory tract infections result in the influx of other immune cells. SARS-CoV-2 infection moderately increased the proportions of myeloid cells in the BAL 3 dpi, with most returning to normal by 9 dpi (Fig S13a). There was no effect of age (Fig S13b). The myeloid influx included cells phenotyped as interstitial macrophages (IMs, Fig 6a, e), neutrophils (Fig 6c, g) and plasmacytoid dendritic cells (pDCs, Fig 6d, h). In contrast, the levels of resident AMs in BAL declined significantly at 3 dpi (Fig 6b, f). The increase in IMs, neutrophils and pDCs at 3 dpi was highly correlated with the levels of viral RNA (Fig 6i-j, Fig S13i), while AMs exhibited an opposite trend. The frequency of conventional dendritic cells (cDCs) declined as pDCs increased in BAL (Fig S13c). An increase in the levels of both classical (CD14^+^CD16^-^) (not shown) and intermediate/inflammatory (CD14^+^CD16^+^) monocytes in BAL was also observed at 3 dpi (Fig S13d). The frequency of myeloid subpopulations increased in BAL was generally reduced in blood (Fig S13e-h), with two exceptions - pDCs and CD14^+^CD16^+^ monocytes, which were increased in the blood as well as BAL (Fig S13g-h). Relative to AMs, IMs have a shorter half-life, exhibit continuous turnover, and may help to maintain homeostasis and protect against continuous pathogen exposure from the environment^14^. Increased recruitment of pDCs to the lungs suggests a potentially important feature of protection from advanced COVID-19 disease in the rhesus macaque model since they are a major source of anti-viral Type I interferons such as IFN-*α*, the levels of which were elevated in the BAL within 1-3 dpi (Fig 3b).

**Figure 6.**
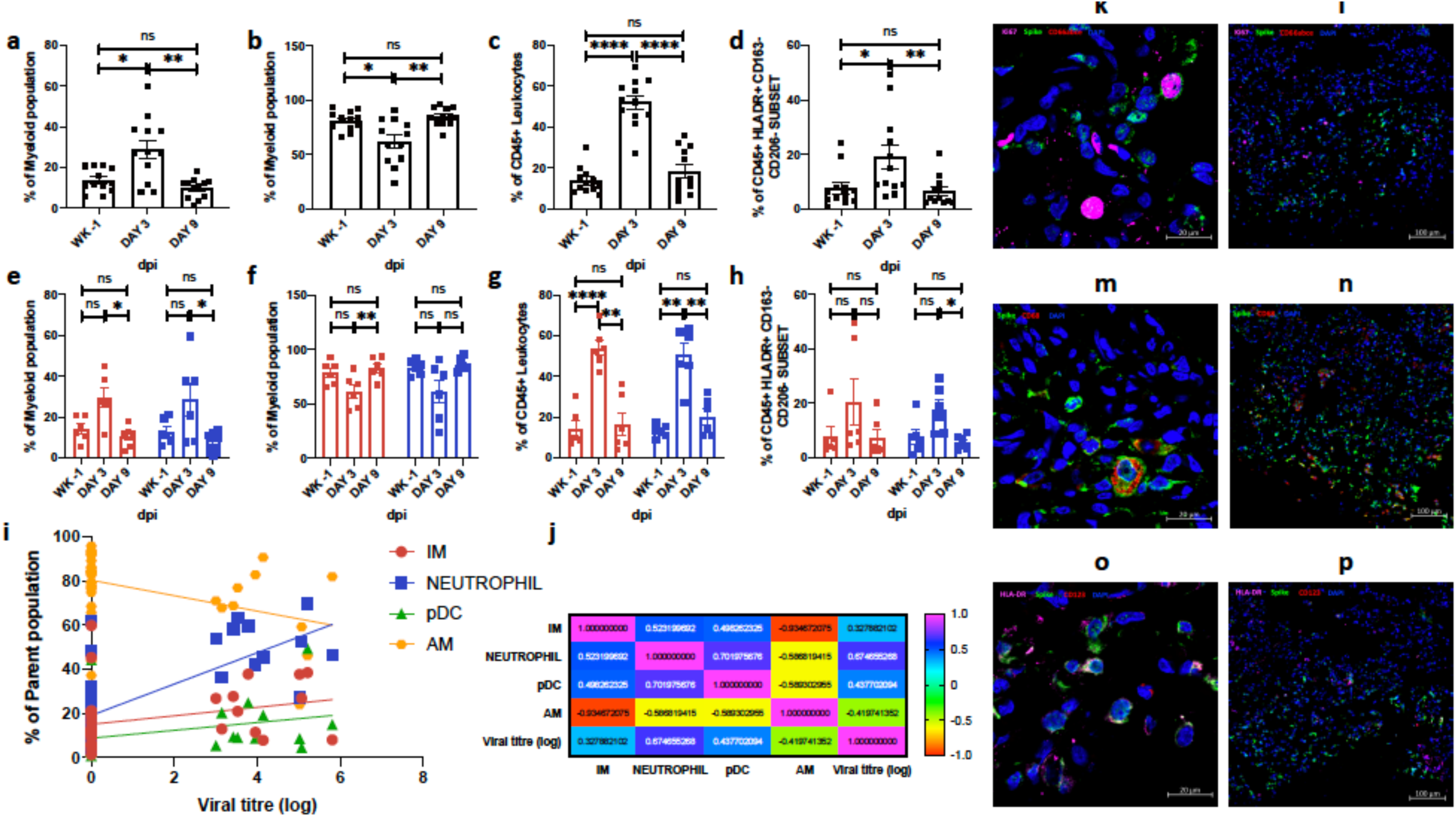
**Longitudinal accumulation of myeloid cells in BAL following SARS-CoV-2 infection in rhesus macaques**. Flow cytometric analysis of BAL IMs (a, e), AMs (b, f), neutrophils (c,g), and pDCs (d, h). Data shown combined for age (a-d) (n=12); data split by age (g-h) (n=6). Data is represented as mean± SEM. (a-d) One way and (e-h) two way Repeated-measures ANOVA with Geisser-Greenhouse correction for sphericity and Tukey’s post hoc correction for multiple-testing (GraphPad Prism 8) was applied. * P<0.005, ** P<0.005, *** P<0.0005. Coloring scheme for e-h – young (blue), old (red). Correlations with Spearman’s rank test between cellular fraction and Log10 viral RNA copy number in BAL (i) and corresponding values for Spearman’s rank correlation coefficient (j) and P value (Suppl. Fig13i). Coloring scheme for i – Neutrophil (blue), IM (red), AM (orange, pDC (green). Multilabel confocal immunofluorescence microscopy of FFPE lung sections from SARS CoV-2 infected Rhesus macaques having a high viral titer at 3 dpi (k-p) with SARS CoV-2 Spike specific antibody (green), KI-67 ^29^, neutrophil marker CD66abce (red) and DAPI (blue) at 10X (k) and 63X (l); SARS CoV-2 Spike (green), pan-macrophage marker CD68 (red) and DAPI (blue) at 10X (m) and 63X (n); SARS CoV-2 Spike (green), HLA-DR ^29^, pDC marker CD123 (red) and DAPI (blue) at 10X (o) and 63X (p).

Multi-label confocal imaging of lung tissues following Ki67 staining depicted that only few of the virally-infected cells in the lung tissue actively proliferated (Fig 5k-p). Detailed analysis of the lung tissue revealed that neutrophils (Fig 6k, l, Fig S14a, d), macrophages (Fig 6m, n, Fig S14b, h) and pDCs (Fig 6o, p, Fig S14c, i) recruited to the lung compartment (Fig 6a-h) harbored high levels of viral proteins (Fig 6k-p, Fig S14). Apart from these, many of the other cell-types contained viral proteins, suggesting a capacity of the virus to infect many cell types and that intact virus may also persist in the lungs. These novel data suggest that rapid influx of specialized subsets of myeloid cells to the lung that are known to express Type I IFNs and other pro-inflammatory cytokines is a key event in the control of SARS-CoV-2 infection.

Infection of macaques also resulted in a significant influx of T cells to the alveolar space by 3 dpi, which normalized by 9 dpi (Fig 7a, b, g). After infection, CD4^+^ T cells expressed significantly lower levels of antigen-experience/tissue residence (CD69; Fig 6c), Th1 (CXCR3; Fig 6d), memory (CCR7; Fig 6f), and activation (HLA-DR) (Fig 6m) markers in BAL. In contrast, the levels of CD4^+^ T cells expressing PD-1 (Fig 6e) and LAG-3 (Fig 6n) were significantly elevated, while those of CD4^+^ T cells expressing CCR5 (Fig 6l) were unchanged. A similar effect was observed in CD8^+^ T cell subsets, where the expression of CD69 (Fig 6h), CXCR3 (Fig 6i), and CCR7 (Fig 6k) was significantly reduced in BAL following infection whereas expression of PD-1 (Fig 6j) and LAG-3 (Fig 6q) in the CD8^+^ T cells was significantly increased. CCR5 (Fig 6o) and HLA-DR (Fig 6p) were unchanged. No differences were observed in T cell responses in young relative to old animals. Taking data from myeloid cells and lymphocytes together, we postulate that the rapid influx of myeloid cells capable of producing high levels of Type I IFNs result in immune control of SARS-CoV-2 infection in macaques, but that this control is not sterilizing. This allows for viral antigens to persist leading T cell recruitment, but with a T cell profile associated with immune modulation and promotion of antigen-mediated T cell anergy/exhaustion (PD-1, LAG3 expression)^15^.

**Figure 7.**
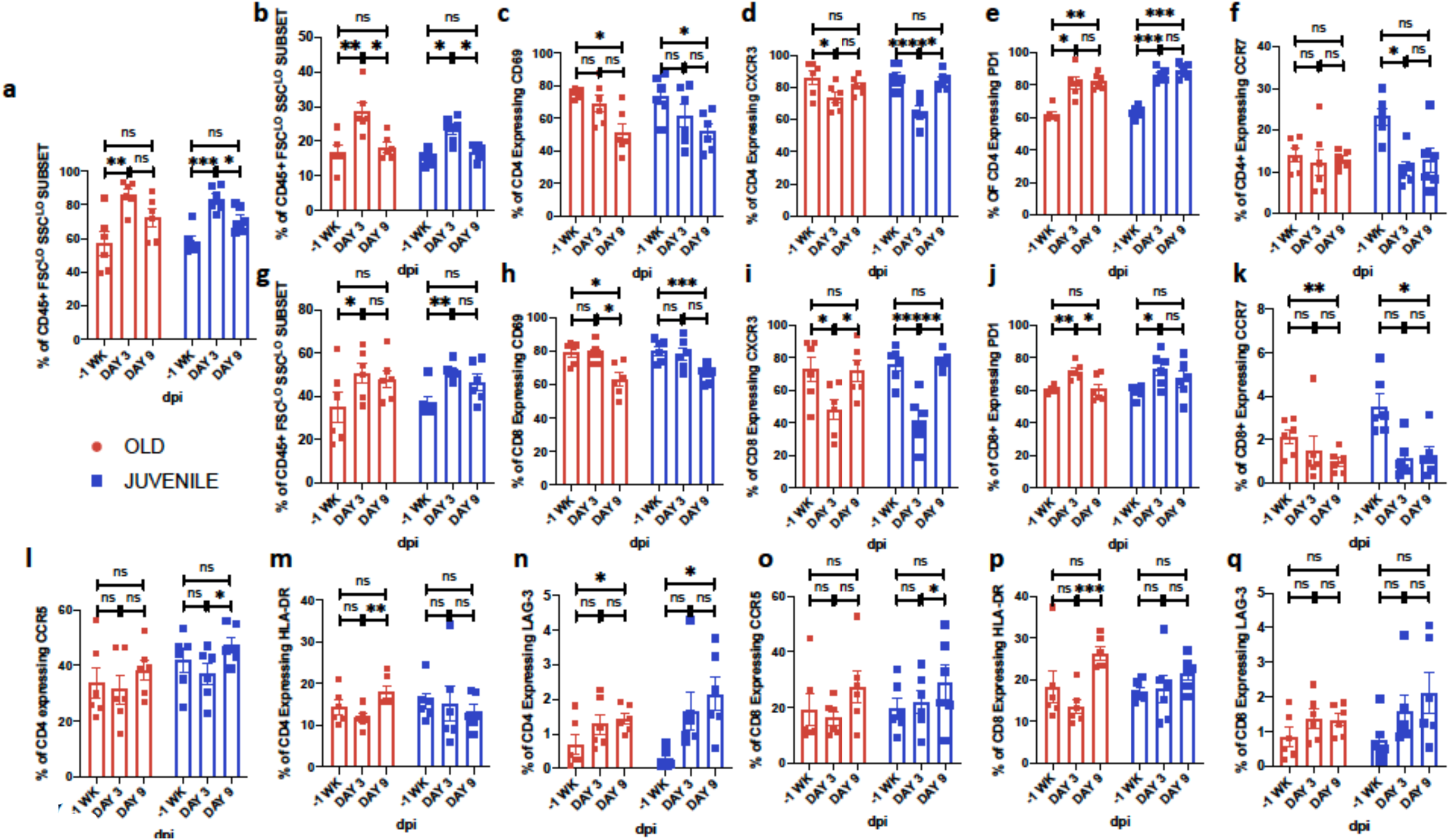
**Longitudinal changes in T cells in BAL following SARS-CoV-2 infection in rhesus macaques**. BAL Frequencies of CD3^+^ T cells (a), CD4^+^ T cells (b), CD8^+^ T cells (g), CD4^+^ T cell subsets expressing early activation marker CD69 (c), CXCR3 (d), PD-1 (e) and memory marker CCR7 (f), CCR5 (l), HLA-DR (m) and LAG-3 (n); CD8^+^ T cell subsets expressing early activation marker CD69 (h), CXCR3 (i), PD-1 (j) and memory marker CCR7 (k), CCR5 (o), HLA-DR (p) and LAG-3 (q). Coloring scheme – young (blue), old (red). Data is represented as mean± SEM. (n=6) Two way Repeated-measures ANOVA with Geisser-Greenhouse correction for sphericity and Tukey’s post hoc correction for multiple-testing (GraphPad Prism 8) was applied. * P<0.005, ** P<0.005, *** P<0.0005.

To extrapolate from phenotype to function, we explored proliferation, immune mediator production, and memory phenotypes. CD4^+^ and CD8^+^ T cells exhibiting proliferative (Fig 8a, g) and memory markers (Fig 8b, h) were significantly increased in BAL after infection whereas CD4^+^ and CD8^+^ T cells expressing naïve (Fig 8c, i) and effector (Fig 8d, j) phenotypes were significantly reduced. The percentage of CD4^+^ (Fig 8e) and CD8^+^ (Fig 8k) T cells expressing IL-2 was significantly elevated in the BAL at 9 dpi. A similar effect was observed for Granzyme-B (GZMB) (Fig 8f, l) which was sustained through 9 dpi. No significant effect of age was observed, although the expression of IL-2 on T cells was higher for young compared old rhesus macaques. Frequencies of CD4^+^ and CD8^+^ expressing interferon-*γ* (IFNG) (Fig S15a, d) and IL-17 (Fig S15b, e) were elevated, but unchanged for TNF-*α* (Fig S15c, f). Greater expression of IFN*γ* was measured on CD4^+^ T cells recruited to the BAL in younger animals, but the differences were not statistically significant. These results suggest that robust cellular immune responses (both CD4^+^ and CD8^+^ T cells) are generated in the lung compartment (BAL) as early as day 3 and maintained at 9 dpi in many instances. Following ex vivo re-stimulation of T cells from BAL at 9 dpi with CoV-specific peptide pools, CD4^+^ T cells expressing IL-2 (Fig 8m), GZMB (Fig 8n), IFN-*γ* (Fig S15g), IL-17 (Fig S15h) and TNF-*α* (Fig S15i) were not statistically elevated beyond baseline values. This was similar for CD8^+^ T cells expressing IL-2 (Fig 8o), GZMB (Fig 8p), IFN-*γ* (Fig S15j), IL-17 (Fig S15k) and TNF-*α* (Fig S15l). In combination with increased expression of the immune-regulatory markers PD-1 and LAG-3, our results suggest that T cells recruited to the lung compartment following SARS-CoV-2 infection are capable of secreting cytokine but fail to generate robust antigen specific responses highlighting the fact that persistent T cell stimulation by viral antigens may generate T cell anergy relatively early in infection and this is promoted by our findings of viral persistence in the respiratory tract.

**Figure 8.**
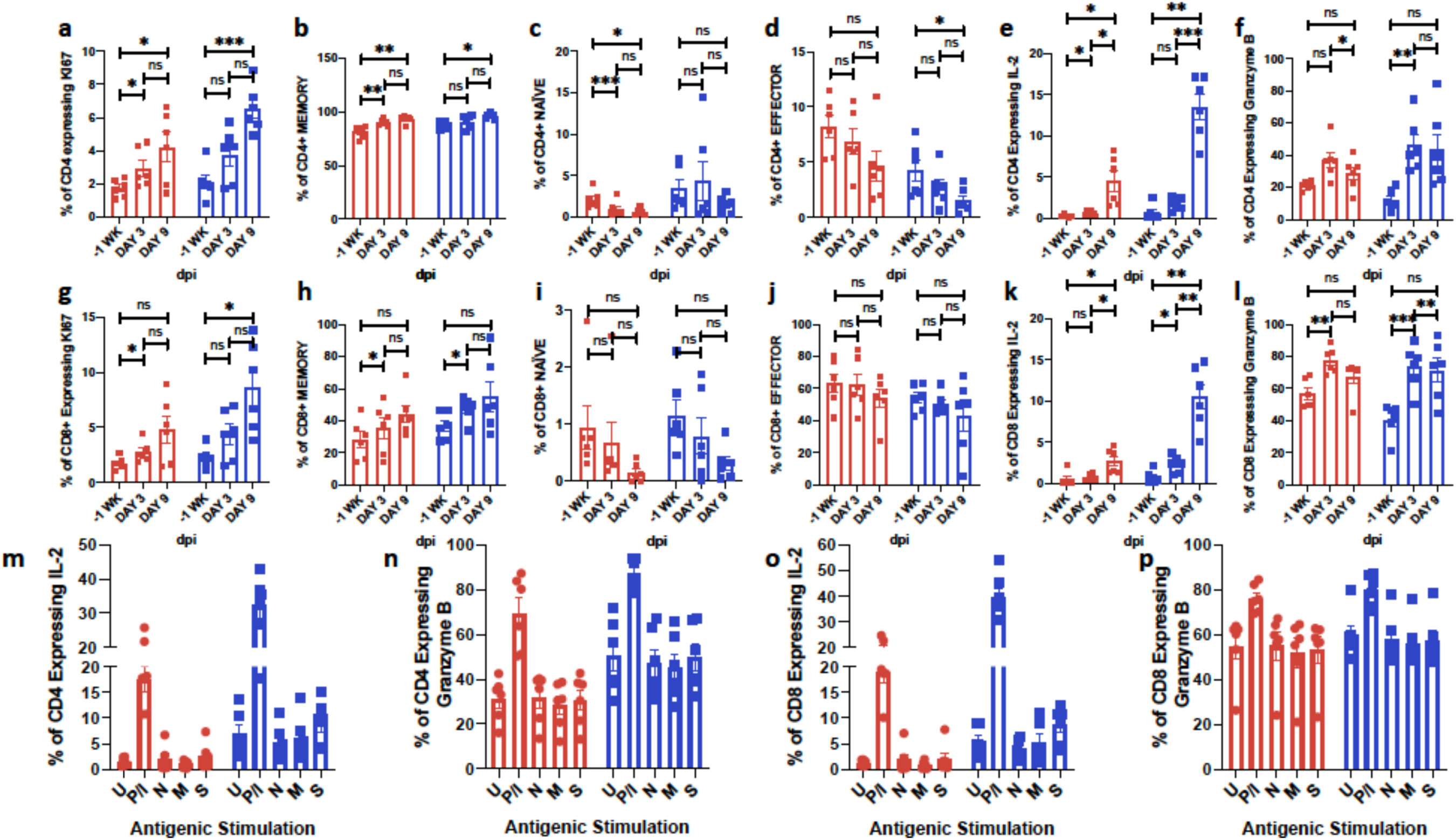
**Longitudinal changes in memory T cells in BAL following SARS-CoV-2 infection in rhesus macaques**. BAL Frequencies of CD4+ T cell subsets expressing KI67 (a), Memory (b), Naïve (c), Effector (d), IL-2 (e) and Granzyme B (f). Frequencies of CD8+ T cell subsets expressing KI67 (g), Memory (h), Naïve (i), Effector (j), IL-2 (k) and Granzyme B (l). BAL cells were stimulated overnight (12-14 hours) with either Mock control (U); PMA-Ionomycin (P/I) or SARS-CoV-2 - specific peptide pools of the nucleocapsid (N), membrane (M) and spike (S) proteins. Antigen specific cytokine secretion in T cells was estimated by flow cytometry. Fraction of CD4^+^ T cells secreting IL-2 (m), Granzyme B (n); CD8+ T cells secreting IL-2 (o) and Granzyme B (p). Coloring scheme – young (blue), old (red). Data is represented as mean± SEM. (n=6) two way Repeated-measures ANOVA with Geisser-Greenhouse correction for sphericity and Tukey’s post hoc correction for multiple-testing (GraphPad Prism 8) was applied. * P<0.005, ** P<0.005, *** P<0.0005.

Immunophenotyping results were confirmed by studying cytokine production in BAL and plasma (Fig S16)^16^. Our results show that the levels of IFN-*α* (Fig S16a), IL-1Ra (Fig S16b), and IL-6 (Fig S16d) were elevated in BAL following infection, but levels rapidly normalized after the 3 dpi peak. Levels of IFN-*α* were also induced in plasma (Fig S16g), but not those of IL-1Ra (Fig S16h), and IL-6 (Fig S16j) or other cytokines studied. Cytokines were not induced at baseline or in procedure control animals. Overall, the longitudinal study results were consistent with the acute infection study in the expression of Type I pro-inflammatory cytokines responsible for viral control (IFN-*α*) and expression of IL-6, which may contribute to a cytokine storm and development of ARDS in a subset of hosts during COVID-19.

Protein levels of ACE-2, one presumed receptor for SARS-CoV-2 in humans, were detected at higher levels in the lungs and nasal epithelia of infected macaques than those in the lungs of naive rhesus macaques (Fig 1j, k). Using RNAseq we also studied if expression of ACE-2 could be detected in macaque lung tissues and elevated in in SARS-CoV-2 infected animals. This was indeed the case (Fig S17a-d) in a statistically significant manner two weeks after infection despite multiple hypothesis correction (Fig S17a, b). Interestingly, ACE-2 expression was significantly higher in young compared to old macaques (Fig S17c, d). These results potentially explain the higher levels of virus that we observed in several samples derived from young macaques in these two cohorts (Fig S8). Expression of transcripts specific for other viral receptors/co-receptors e.g., Cathepsin-L, CD147 or TMPRSS2 was not significantly altered in the lung two weeks after infection (Fig S17a).

Altogether, our results show that rhesus macaques, baboons and marmosets can all be infected with SARS-CoV-2 but exhibit differential progression to COVID-19. While marmosets exhibit mild infection, macaques are characterized by the presence of moderate progressive pneumonia that is rapidly resolved. This is accompanied by a marked reduction in lung and nasal viral loads. Baboons appear to have the most lung pathology, and the level of viral shedding and persistence in extra-respiratory compartment is also greater in this model. Furthermore, we show the importance of state-of-the-art, non-invasive imaging – cone beam CT scanning, and the application of innovative algorithms to identify the extent of lung involved in pneumonia, in developing models of COVID-19. This provided us with a quantifiable metric that lent itself to accurately assessing the efficacy of vaccines or the impact of therapeutic interventions.

Our results also point out, for the first time, that SARS-CoV-2 infection is associated with dynamic influxes of specific subsets of myeloid cells to the lung, particularly IMs, neutrophils and pDCs, and that viral proteins can be detected in these cells. These cellular influxes are likely due to a strong viral-induced myelopoeisis. This may help explain both development of COVID-19 pneumonia and subsequent control via expression of a strong Type I IFN response and expression of other pro-inflammatory cytokines. We speculate that these responses clear the majority of virus, and, in doing so, lead to eventual resolution of pneumonia, while limiting a progressive cytokine storm and ARDS in the majority of hosts. Macaques have served as excellent models of infectious diseases and vaccine development efforts^17–19^, and this model permits lung imaging and detailed immune evaluations. Given the ability to reproducibly measure viral loads in NS and BAL, and quantify lung involvement by CT scans and hyperdensity analyses, we expect this model to play a critical role in the preclinical testing of novel candidate vaccines against SARS-CoV-2 infection and/or COVID-19 disease in development. Experiments in rhesus macaques can also evaluate safety and immunogenicity, including the important issue of antibody-mediated immune enhancement. Since mild-to-moderate COVID-19 disease that follows SARS-CoV-2 infection in rhesus macaques is short-lived, it follows that vaccine safety and efficacy studies can be evaluated in short term studies.

However, detection of both virus and its protein antigens over two weeks in macaques, baboons and even marmosets, indicates viral persistence rather than sterilizing immunity. Support for this comes from the finding of PD-1 and LAG-3 expression by CD4^+^ and CD8^+^ T cells in the lung and lack of induction of antigen-specific immune effector cytokine production by these cells. Characterization of these responses is particularly important considering that T cell responses, particularly T helper responses, play key roles in shaping the nature of downstream B cell responses and production of antibodies. It is likely that in immunocompromised patients, persistent presence of SARS-CoV-2 could lead to exacerbated disease. Since COVID-19 has disproportionately affected the aging human population, we included age as an independent variable in our studies. Although there were several smaller changes observed in older animals, old and young animals both resolved infection. While it is possible that NHPs do not completely model all aspects of COVID-19 in humans, these findings suggest that underlying conditions which impact immunity such as defined and undefined co-morbidities, rather than aging per se, may be responsible for the greater morbidity and mortality observed due to COVID-19 in the aged human population (and a subset of younger individuals). Baboons developed more extensive disease and pathology with more widespread and severe inflammatory lesions compared to rhesus macaques. Baboons are also a preferred model of cardiovascular and metabolic diseases including diabetes ^20–22^, and therefore further development of the baboon model may prove especially useful for the study of co-morbidities with COVID-19 such as diabetes, cardiovascular disease, and aging.

## Methods

### Study approval

All of the infected animals were housed in Animal Biosafety Level 3 or 4 (ABSL3, ABSL4) at the Southwest National Primate Research Center where they were treated per the standards recommended by AAALAC International and the NIH Guide for the Care and Use of Laboratory Animals. Sham controls were housed in ABSL2. The animal studies in each of the species were approved by the Animal Care and Use Committee of the Texas Biomedical Research Institute and as an omnibus Biosafety Committee protocol.

### Animal studies and clinical evaluations

16 (eight young and eight young, see Table S1 for details) Indian-origin rhesus macaques *(Macaca mulatta)*, and six African-origin baboons (*Papio hamadryas*) all from SNPRC breeding colonies, were exposed via multiple routes (ocular, 100 μL; intranasal, 200 μL - using a Teleflex Intranasal Mucosal Atomization Device; intratracheal, 200 μL - using a Teleflex Laryngo-Tracheal Mucosal Atomization Device) of inoculation to 500 μL of an undiluted stock of SARS-CoV-2, which had a titer of 2.1E+06 pfu/mL, resulting in the administration of 1.05×106 pfu SARS-CoV-2. SARS-CoV-2 generated from isolate USA-WA1/2020 was used for animal exposures. A fourth cell-culture passage (P4) of SARS-CoV-2 was obtained from Biodefense and Emerging Infections Research Resources Repository (BEI Resources, catalog number NR-52281, GenBank accession number MN985325.1) and propagated at Texas Biomed. The stock virus was passaged for a fifth time in Vero E6 cells at a multiplicity of infection (MOI) of approximately 0.001. This master stock was used to generate a sixth cell culture passage exposure stock by infecting VeroE6 cells at a MOI of 0.02. The resulting stock had a titer of 2.10 x 106 PFU/mL and was attributed the Lot No. 20200320. The exposure stock has been confirmed to be SARS-CoV-2 by deep sequencing and was identical to published sequence (MN985325). strain USA-WA1/2020 (BEI Resources, NR-52281, Manassas, VA). Six Brazilian-origin common marmosets (*Callithrix jacchus*) were also infected via the combined routes (80µL intranasal; 40µL ocular [20µL/eye]; 40µL oral performed twice for a total of 160µL intranasal, 80µL ocular; 80µL oral and 100µL IT) of the same stock. The total target dose presented to marmosets was 8.82E+05 pfu/mL. Four macaques, baboons and marmosets each were sham-infected with DMEM-10 media (the storage vehicle of the virus), to be used as procedural controls. Infected animals were euthanized for tissue collection at necropsy, and control animals were returned to the colony. Macaques were enrolled from a specific pathogen-free colony maintained at the SNPRC and were tested free from SPF-4 (simian retrovirus D, SIV, STLV-1 and herpes B virus). All animals including the baboons and the marmosets were also free of *Mycobacterium tuberculosis*. Animals were monitored regularly by a board-certified veterinary clinician for rectal body temperature, weight and physical examination. Collection of blood, BAL, nasal swab, and urine, under tiletamine-zolazepam (Telazol) anesthesia was performed as described (Table S1), except that BAL was not performed in marmosets. Four macaques were sampled daily until euthanized at 3dpi. All other macaques and all the baboons were sampled at 0, 3, 6, 9, 12 dpi and at euthanasia (BAL performed weekly). Blood was collected for complete blood cell analysis and specialized serum chemistries. Animals were observed daily to record alert clinical measurements. Nasal (longitudinal) or nasopharyngeal (acute) swabs and BALs were obtained to measure viral loads in a longitudinal manner, as described earlier ^11^. Briefly, in a sitting position, the larynx was visualized and a sterile feeding tube inserted into the trachea and advanced until met with resistance. Up to 80ml of warm sterile saline was instilled, divided into multiple aliquots. Fluid was aspirated and collected for analysis.

### Chest X-Rays

Clinical radiographic evaluation was performed as following: The lungs of all animals were imaged by conventional (chest radiography, CXR), as previously described ^23^. Three view thoracic radiographs (ventrodorsal, right and left lateral) were performed at all sampling time points. High-resolution computed tomography (CT) was performed daily through 3 dpi in 4 infected macaques and on 6 and 12 dpi in 3 young and 3 old macaques as described in the next section. Images were evaluated by a board-certified veterinary radiologist and scored as normal, mild moderate or severe disease. The changes were characterized as to location (lung lobe) and distribution (perivascular/peribronchial, hilar, peripheral, diffuse, multifocal/patchy).

### CT Imaging and quantitative analysis of lung pathology

The animals were anesthetized using Telazol (2-6mg/kg) and maintained by inhaled isoflurane delivered through Hallowell 2002 ventilator anesthesia system (Hallowell, Pittsfield, MA). Animals were intubated to perform end-inspiratory breath-hold using a remote breath-hold switch. Lung field CT images were acquired using Multiscan LFER150 PET/CT (MEDISO Inc., Budapest, Hungary) scanner. Image analysis was performed using 3D ROI tools available in Vivoquant (Invicro, Boston, MA). Percent change in lung hyperdensity was calculated to quantify lung pathology (1, 2). The lung volume involved in pneumonia, was quantified as follows: briefly, lung segmentation was performed using a connected thresholding feature, to identify lung ROI by classifying all the input voxels of scan in the range of −850 HU to −500 HU. Smoothing filters were used to reassign every ROI voxel value to the mode of the surrounding region with defined voxel radius and iterations to reconstruct the Lung ROI. Thereafter, global thresholding was applied to classify the voxels within Lung ROI in the range of −490 HU to +500 HU to obtain Lung hyperdensity ROI. The resultant ROIs were then rendered in the maximum intensity projection view using the VTK feature.

### Viral RNA determination

Viral RNA from plasma/sera, BAL, urine, saliva, and swabs (nasal/nasopharyngeal, oropharyngeal, rectal) and lung homogenates was determined by RT-qPCR and viral RNA isolation as previously described for MERS-CoV and SARS-CoV (12, 27, 28). RNA extraction from fluids was performed using the EpMotion M5073c Liquid Handler (Eppendorf) and the NucleoMag Pathogen kit (Macherey-Nagel). 100 µL of test sample were mixed with 150 µL of 1X DPBS (Gibco) and 750 µL TRIzol LS. Inactivation controls were prepared with each batch of samples to ensure no cross contamination occurred during inactivation. Samples were thawed at room temperature and then, for serum, swabs and urine samples 10µg yeast tRNA was added, along with 1 x 103 pfu of MS2 phage (*Escherichia coli* bacteriophage MS2, ATCC). DNA LoBind Tubes (Eppendorf) were prepared with 20 µL of NucleoMag B-Beads (NucleoMag Pathogen kit, Macherey-Nagel) and 975 µL of Buffer NPB2 (NucleoMag Pathogen kit, Macherey-Nagel). After centrifugation, the upper aqueous phase of each sample was transferred to the corresponding new tube containing NucleoMag B-Beads and Buffer NPB2. The samples were mixed using HulaMixer (Thermo Fisher Scientific Inc.) rotating for 10 min at room temperature. Samples were then transferred to the sample rack on EpMotion M5073c Liquid Handler (Eppendorf) for further processing according to NucleoMag Pathogen kit instructions. For viral RNA determination from tissues, 100mg of tissue was homogenized in 1mL Trizol Reagent (Invitrogen, Grand Island, NY, USA) with a Qiagen (Germantown, MD, USA) steel bead and Qiagen Stratagene TissueLyser. For detection of infectious virus, briefly, tissues were homogenized 10% w/v in viral transport medium using Polytron PT2100 tissue grinders (Kinematica). After low-speed centrifugation, the homogenates were frozen at −70°C until they were inoculated on Vero E6 cell cultures in 10-fold serial dilutions. The SARS-CoV-2 RT-qPCR was performed using a CDC-developed 2019-nCoV_N1 assay with the TaqPath™ 1-Step RT-qPCR Master Mix, CG (ThermoFisher). The assays were performed on a QuantStudio 3 instrument (Applied Biosystems) with the following cycling parameters: Hold stage 2 min at 25°C, 15 min at 50°C, 2 min at 95°C. PCR stage 45 cycles of 3 s at 95°C, 30 s at 60°C. Primer and probe info: 2019-nCoV_N1-F: GACCCCAAAATCAGCGAAAT (500nM); 2019-nCoV_N1-R: TCTGGTTACTGCCAGTTGAATCTG (500 nM); 2019-nCoV_N1-P FAM/MGB probe: ACCCCGCATTACGTTTGGTGGACC (125nM).

### Pathology

Animals were euthanized and complete necropsy was performed. Gross images (lung, spleen, liver) and organ weights (lymph nodes, tonsil, spleen, lung, liver, adrenal glands) were obtained at necropsy. Representative samples of lung lymph nodes (inguinal, axillary, mandibular and mediastinal), tonsil, thyroid gland, trachea, heart, spleen, liver, kidney, adrenal gland, digestive system (stomach, duodenum, jejunum, ileum, colon, and rectum), testes or ovary, brain, eye, nasal tissue, and skin were collected for all animals. Tissues were fixed in 10% neutral buffered formalin, processed to paraffin, sectioned at 5 um thickness, stained with hematoxylin and eosin utilizing standard methods, and evaluated by a board-certified veterinary pathologist.

### Tissue processing, flow cytometry, multiplex cytokine analyses, immunohistochemistry, multicolor confocal microscopy and RNAseq for immune evaluations

Flow cytometry was performed as previously described ^24–26^ on blood and BAL samples collected on time points days 3, 6, 9, 12, and at endpoint, which occurred at 14-17 dpi for various animals. A comprehensive list of antibodies used in these experiments is provided in Table S7. For evaluations on peripheral blood, PBMC were prepared as previously described. Briefly, Cellular phenotypes were studied using antibodies: CD3 (clone SP34-2), CD4 (clone L200), CD69 (clone FN50), CD20 (2H7), CD95 (clone DX2), KI67 (B56), CCR5 (3A9), CCR7(clone 3D12), CD28 (clone CD28.2), CD45 (clone D058-1283), CXCR3 (clone 1C6/CXCR3), HLA-DR (clone L243), CCR6 (clone 11A9), LAG-3 (Polyclonal, R&D Systems, Minneapolis, MN, USA), CD123 (clone 7G3), CD14 (clone M5E2), CD206 (clone 206), CD16 (clone 3G8), CD163 (GHI/61), CD66abce (Clone TET2, Miltenyi Biotech, USA), CD40 (clone 5C3), IL-2(clone MQ1-17H12), Granzyme-B (clone GB11) all purchased from BD Biosciences (San Jose, CA, USA) unless specified. CD8 (clone RPA-T8), CD11c (clone 3.9), TNF-alpha (clone MAb11), IFN-gamma (clone B27), IL-17 (clone BL168) and PD-1 (clone EH12.2H7) were purchased from BioLegend, San Diego, CA, US. For antigenic stimulation cells were cultured overnight with SARS-CoV-2 specific peptide pools of the nucleocapsid (N), membrane (M) and spike (S) proteins (PepTivator SARS-CoV-2 peptide pool, Miltenyi Biotech, USA). A detailed gating strategy for detection and enumeration of various cellular phenotypes is described (Fig S18).

Immuno-histochemistry was performed on 4 μm thick sections of lung, nasal cavity and tonsils. The sections were baked at 65°C for 30 min followed by de-paraffinization using Xylene and subsequent hydration with decreasing gradations of ethanol as described ^11, 27^. Heat induced antigen retrieval was performed using Sodium citrate buffer (10mM, pH 6.0) followed by blocking (3 % BSA in TBST for 1 h at 37°C). For SARS CoV-2 detection, specimens were incubated with rabbit SARS CoV-2 spike (S) antibody (ProSci, USA, 1:200, 37°C for 2 h) or anti-SARS CoV-2 nucleocapsid (N) antibody (Sino Biologicals, USA, 1:100, 2h at 37°C). Antihuman ACE-2 (R&D Systems, USA, 1:50, 2h at 37°C) was used for identification of ACE-2. Mouse anti-human CD66abce-PE conjugated (Miltenyi Biotech, USA, 1:20, 2 h at 37°C) was used for identification of neutrophils; mouse CD68 (Thermo Fisher Scientific, USA, 1:100, 2 h at 37°C) for macrophages and pDC’s were identified by co-staining of PE conjugated mouse anti-human CD123 (BD Biosciences, USA, 1:20, 37°C for 2h) and mouse anti-human HLA-DR antibody (Thermo Fisher Scientific, USA, 1:100, 2 h at 37°C). Also, mouse anti-Ki67 (BD Biosciences, USA, 1:50, 2 h at 37°C) was used for detection of actively proliferating cells. Chicken anti-rabbit IgG (H+L), Alexa Fluor 488 conjugate; goat anti-mouse IgG (H+L), Alexa Fluor 647 conjugate; donkey anti-mouse IgG (H+L), Alexa-Fluor 555 conjugate secondary antibodies (Thermo Fisher Scientific, USA, 1:400, 1 h at 37°C) were used for labelling Spike, Ki67 and HLA-DR, CD68 primary antibodies respectively. Tissue sections were then stained with DAPI (Thermo Fisher Scientific, USA, 1:5000, 5 min at 37°C) with subsequent mounting with Prolong Diamond Antifade mountant (Thermo Fisher Scientific, USA). Ziess LSM 800 confocal microscope was used to visualize the stained sections (10X, 20X and 63X magnification).

RNA was isolated, RNAseq performed and data analyzed as described ^16^.

### Statistical analyses. Statistical analyses

Graphs were prepared and statistical comparisons applied using GraphPad Prism version 8 (La Jolla, CA). Various statistical comparisons were performed viz. 2-tailed Student’s t-test, ordinary analysis of variance (ANOVA) or one-way or two-way repeated measure analysis of variance (rmANOVA) with Geisser-Greenhouse correction for sphericity and Tukey’s post hoc correction for multiple-testing (GraphPad Prism 8) was applied wherever applicable and as described in the figure legends. For Correlation analysis, Spearman’s rank test was applied. Statistical differences between groups were reported significant when the p-value is less than or equal to 0.05. The data are presented in mean ± SEM.

## Author Contributions

DK, LSS, RC, LDG designed these studies. DKS, SRG, BS, JC, KJA, RE, T-HL, MG, YG-G, RS, AC, RT, MG, CA, AB, JF, CB, HS, LP, JC, AM, BK, RNP, PE, VH, XA, AB, CK, MA, BR conducted the experiments and acquired the data. EC, AG, JD, SH-U, PAF, CNR, KS, CC, CH, OG, JD, AKV, CH, EJD and KB provided veterinary, veterinary pathology, imaging, colony management or management expertise; DKS, SRG, BS, KJA, AC, MG, EC, RNP, JS, AO, DKA, RC, BR, TJCA, SAK, MM, LDG, RC and DK analyzed the data; DK wrote the paper; LSS, JT, LDG, RC, JBT, KB, EC, LMS, JLP, SG and DKS provided assistance with writing the paper.

## Acknowledgments

We acknowledge exceptional work by our SNPRC veterinary technical and care staff (especially the veterinary technical/animal care groups headed by Tyneshia Camp, Wade Hodgkins, Manuel Aguilar, David Vandenberg and Laura Rumpf) as well as the entire SNPRC and Texas Biomed administrative staff, especially Helen Hawn, for assistance with this study, especially during trying times.

---

**Figure S1.**
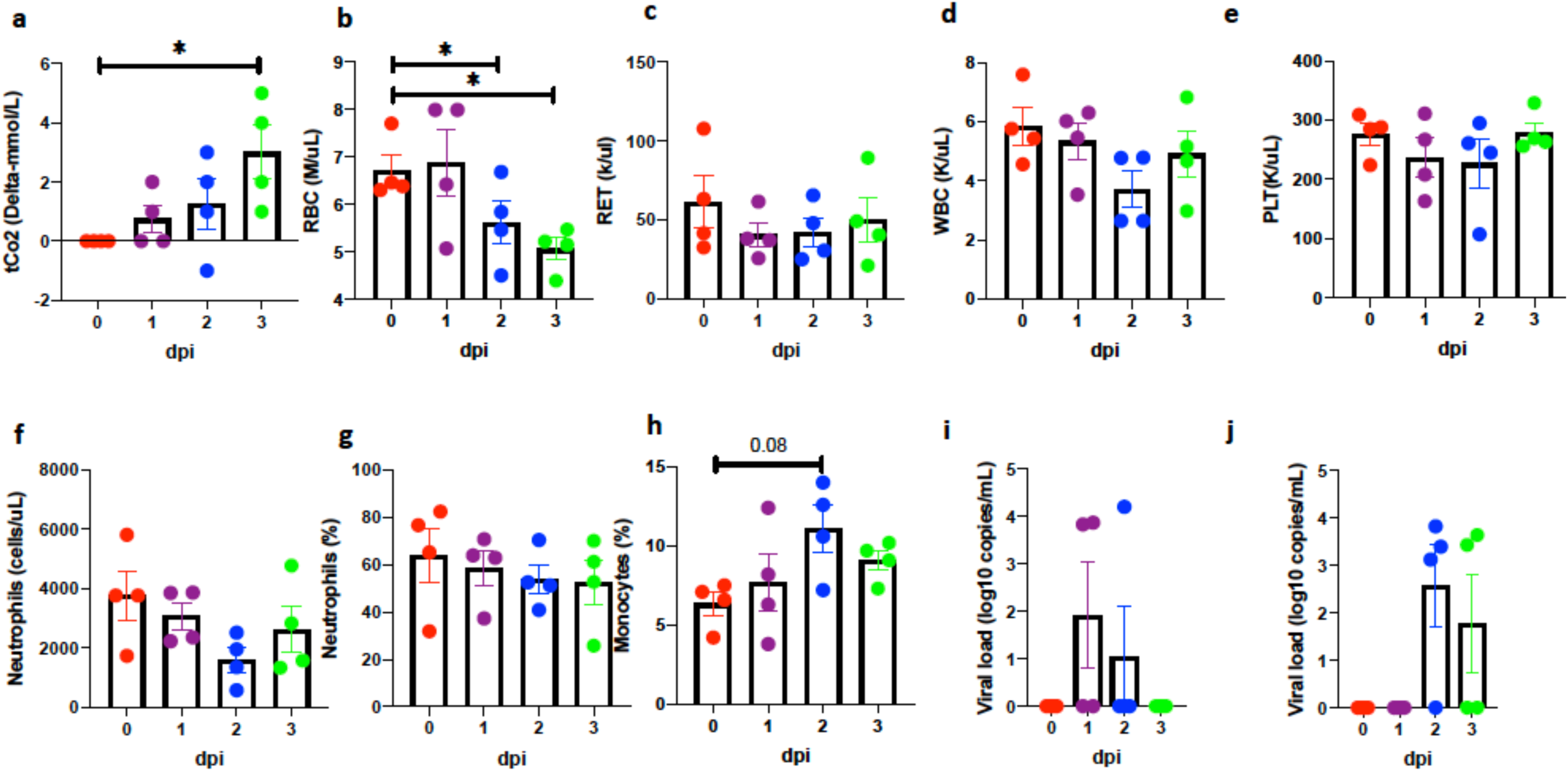
**Clinical correlates in short-term (0-3 dpi) rhesus macaques**. Serum levels of tCO2 (D-mmol/L) (a), and whole blood levels of Red Blood Cells (RBCs) (million/mL) (b), reticulocytes (K/mL) (c), White Blood Cells (WBCs) (K/mL) (d), platelets (K/uL) (e), Neutrophils (K/mL) (f), percentage of Neutrophils (g), percentage of monocytes (h). Viral RNA (log_10_ copies/mL were measured by RT-PCR in saliva (i), and rectal swab (j).) Data is represented as mean± SEM (n=4). One way Repeated-measures ANOVA with Geisser-Greenhouse correction for sphericity and Tukey’s post hoc correction for multiple-testing (GraphPad Prism 8) was applied. * P<0.005, ** P<0.005, *** P<0.0005.

**Figure S2.**
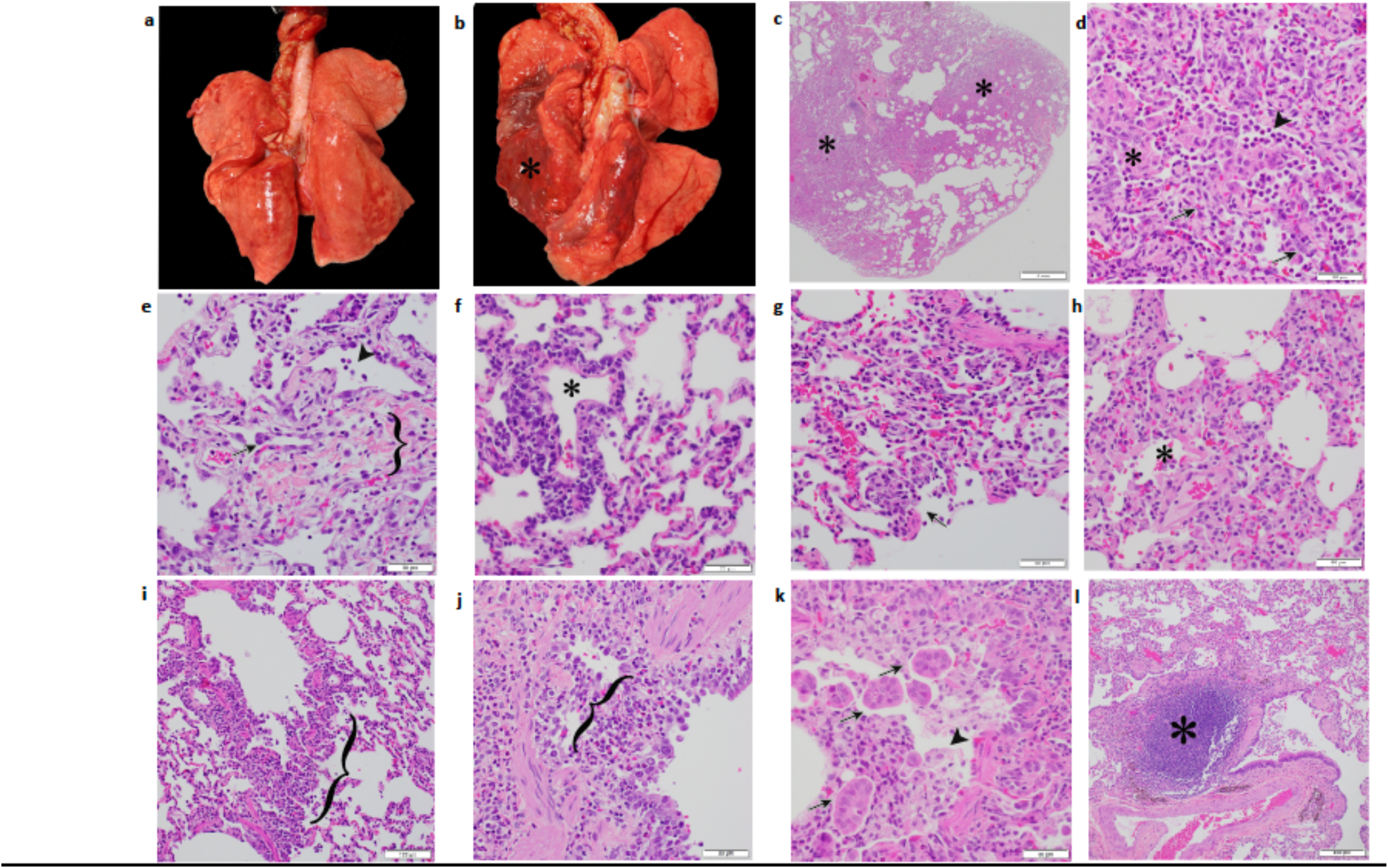
Gross and histopathologic findings of young and aged male and female Rhesus macaques experimentally exposed to COVID19 - 3 dpi. Young male Rhesus macaque. Lung was grossly unremarkable (a). Aged male Rhesus macaque. Lung. The dorsal aspect of the lungs was mottled red (*) (b). Aged male Rhesus macaque. Lung. Sub gross image showing extensive areas of consolidation (*) (c). Aged male Rhesus macaque. Lung. Moderate interstitial pneumonia with scattered type II pneumocytes (arrow), neutrophils (arrowhead), and intra-alveolar fibrin deposition (*) (d). Aged female Rhesus macaque. Lung. Mild interstitial pneumonia with scattered syncytial cells (arrow), neutrophils (arrowhead), and expansion of alveolar walls by fibrosis (bracket) (e). Young female Rhesus macaque. Lung. Vasculitis. Vascular wall disrupted by infiltrates of mononuclear cells and lesser neutrophils. Vessel lumen marked by (*) (f). Young female Rhesus macaque. Lung. Mild interstitial pneumonia. Alveolar spaces contain neutrophils and cellular debris (necrosis, arrow) (g). Young female Rhesus macaque. Lung. Mild interstitial pneumonia. Alveolar spaces (*) contain neutrophils and eosinophilic fluid (edema) (h). Young female Rhesus macaque. Lung. Bronchiolitis. Bronchiolar wall expanded by infiltrates of lymphocytes and macrophages (bracket) (i). Young male Rhesus macaque. Lung. Bronchitis. Bronchial wall expanded by infiltrates of eosinophils that expand and disrupt the epithelium and smooth muscle (bracket) (j). Young female Rhesus macaque. Lung. Bronchitis. Bronchial lumen contains macrophages (arrowhead), cellular debris, and syncytial cells (arrow) (k). Aged female Rhesus macaque. Lung. Area of bronchiolar associated lymphoid tissue (BALT) (*) (l). All slides were stained with H&E.

**Fig S3.**
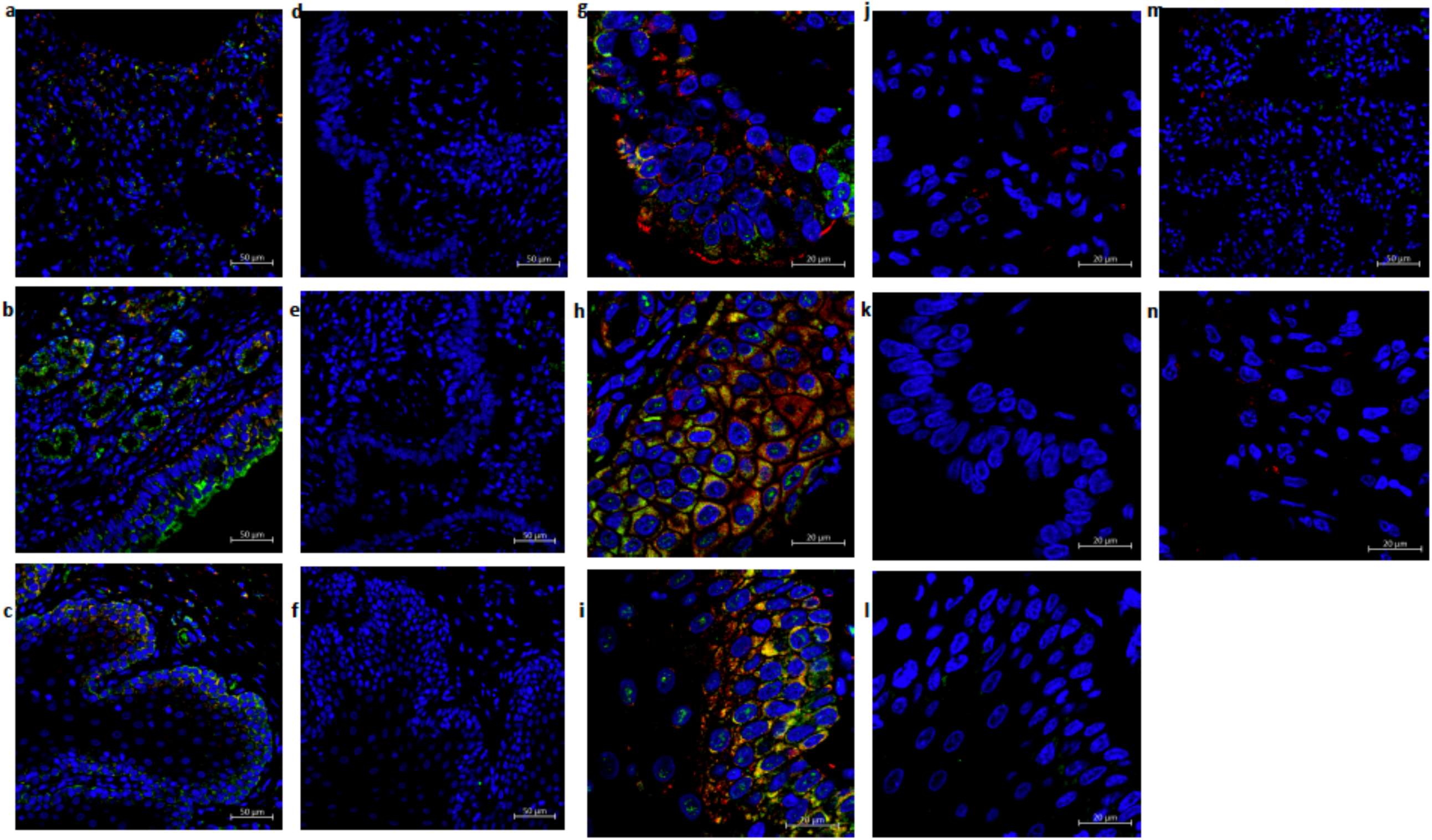
Multi-label confocal immunofluorescence microscopy of lungs (20X-a, 63X-g), nasal epithelium (20X-b, 63x-h) and tonsil (20X-c,63X-i) with SARS CoV-2 N specific antibody (green), DAPI (blue) and ACE-2 (red). Rabbit IgG isotype control antibody was used to rule out non-specific staining in lungs (20X-d, 63X-j), nasal epithelium (20X-e, 63x-k) and tonsil (20X-f, 63X-l). Staining in naïve rhesus macaque lung tissues did not show N signal in lungs (m) or nasal epithelium (n).

**Figure S4.**
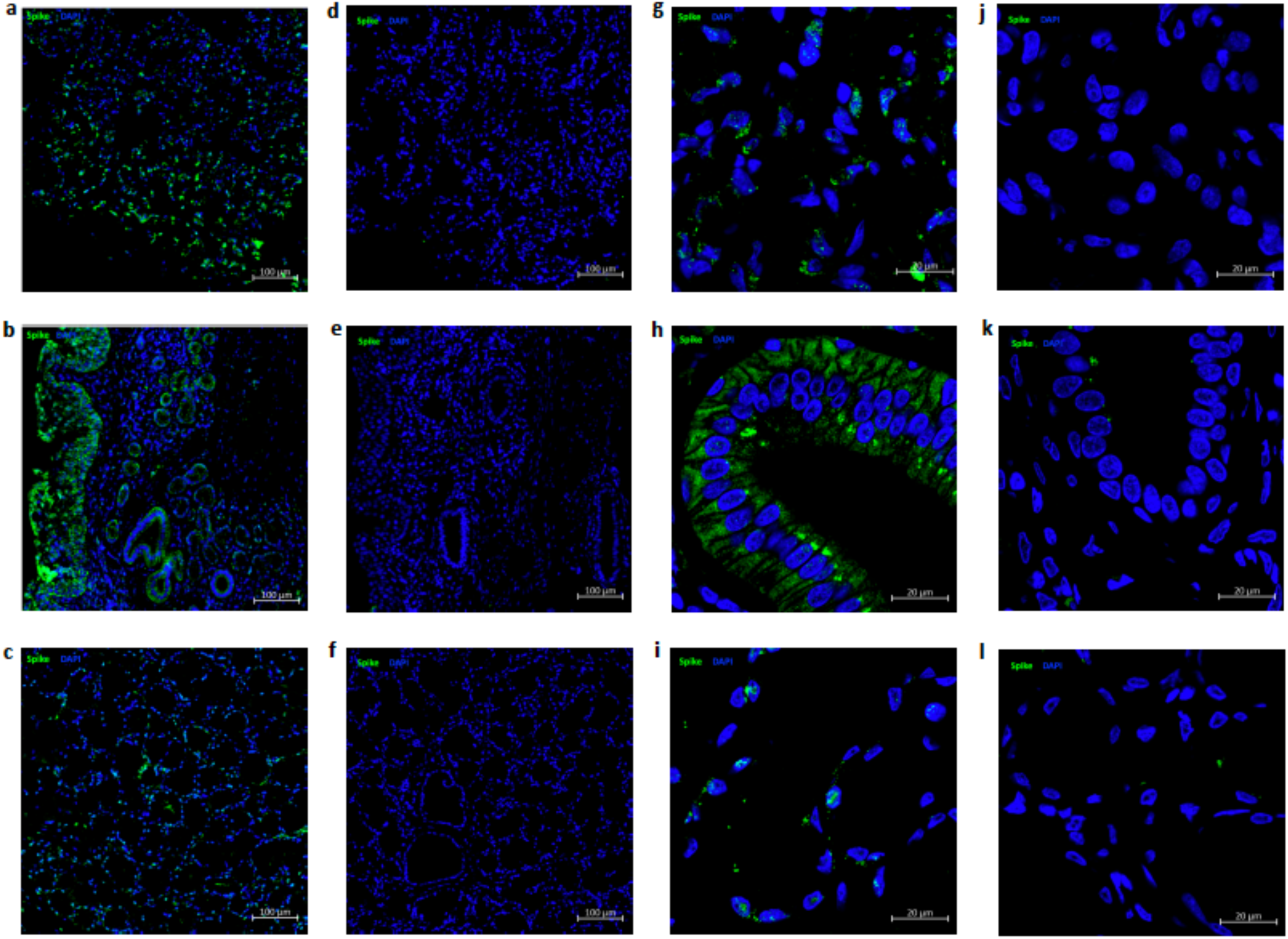
Multi-label confocal immunofluorescence microscopy of lungs (10X-a, 63X-g), nasal epithelium (10X-b, 63x-h) and tonsil (10X-c,63X-i) with SARS CoV-2 S specific antibody (green) and DAPI (blue). Rabbit IgG isotype control antibody was used to stain the tissues to rule out any non-specific staining. The panels showing isotype control staining include: lungs (10X-d, 63X-j), nasal epithelium (10X-e, 63X-k) and tonsil (10X-f, 63X-l).

**Figure S5.**
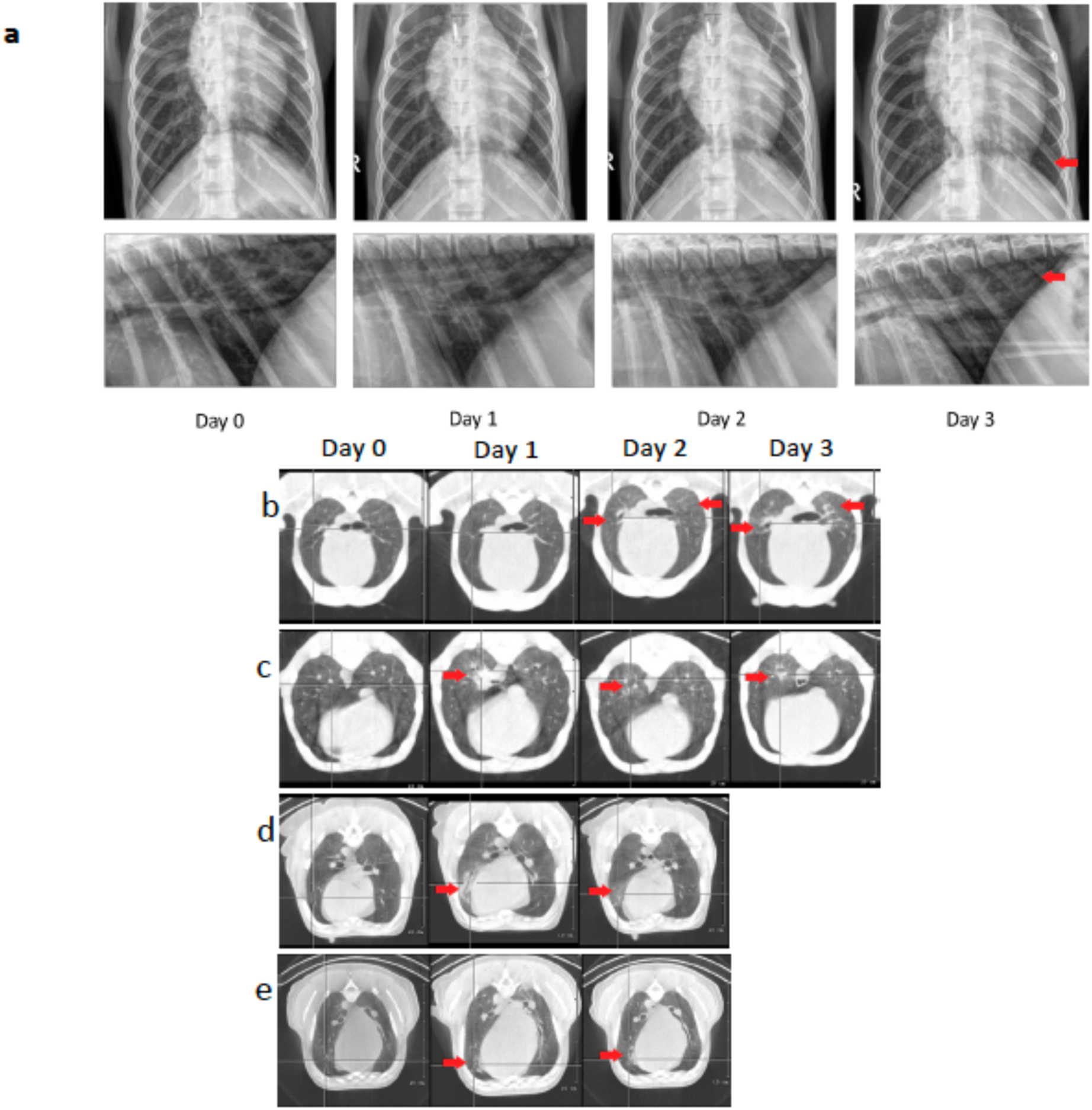
**Radiology of Rhesus macaques experimentally exposed to COVID19 - 3 dpi**. CXR Radiographs showing ventrodorsal and right lateral views(a). Day 0: Normal, Day 1: Mild left caudal interstitial opacity with minimal diffuse right interstitial opacity, Day 2: Mild multifocal interstitial pattern (red arrow), Day 3: Mild multifocal interstitial pattern with patchy region in left caudal lobe (red arrow). CT scan axial view showing lesion characteristics in rhesus macaques infected with SARS-CoV-2 (b) at baseline and Day 1-3 dpi. As seen in (b) ground glass opacity seen on Day 2 dpi intensified on Day 3 dpi. (c) and (d) show lesions that appear on Day 1 show gradual resolution on Day 2-3 dpi whereas lesion in panel (e) observed on Day 1 dpi showed only minimal changes on Day 2. Red arrow point towards lung lesions with high attenuation.

**Figure S6.**
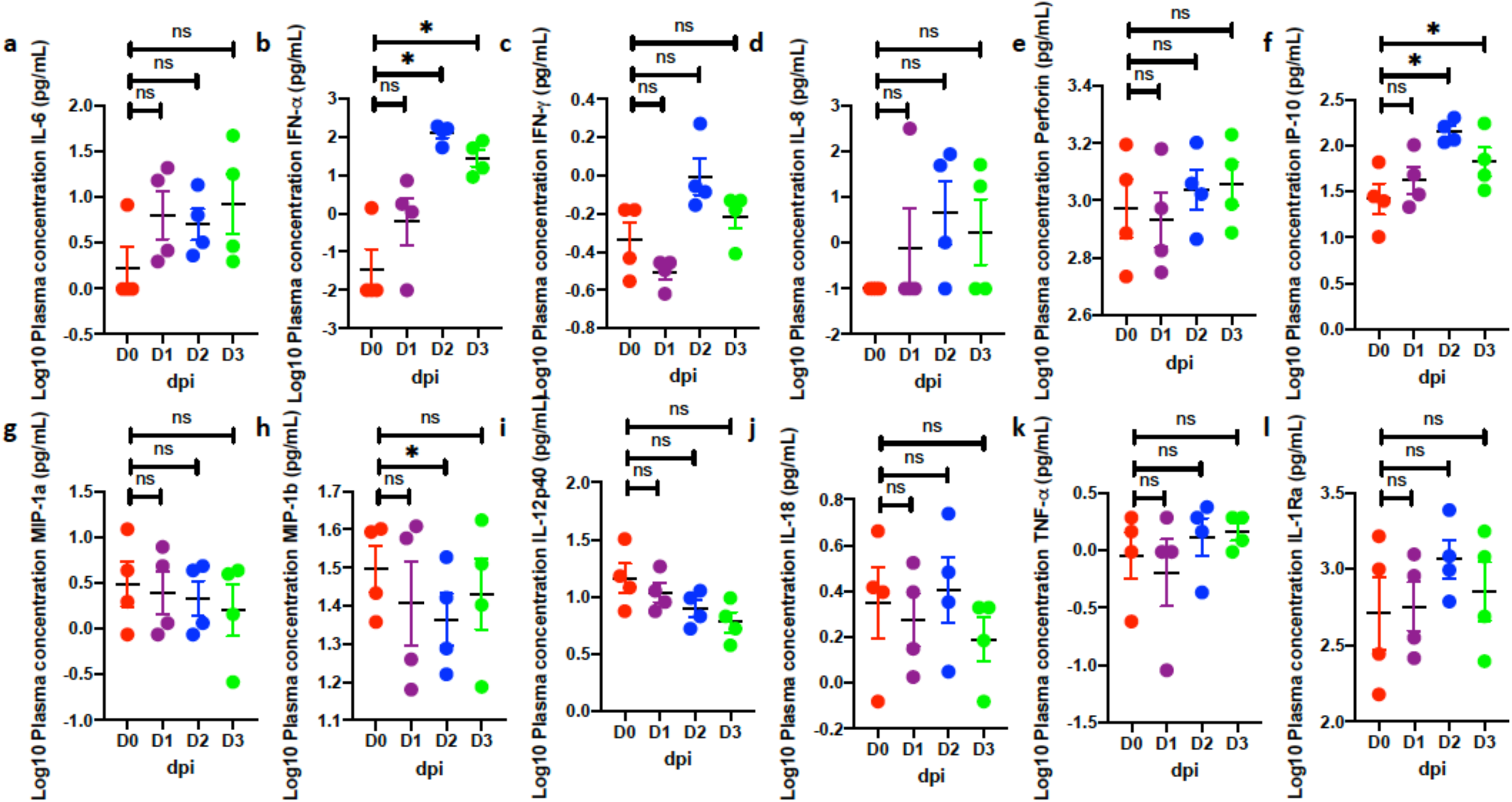
**SARS-CoV-2 induced cytokines in plasma**. Simultaneous analysis of multiple cytokines by Luminex technology in the plasma of rhesus macaques over 0-3 dpi. Levels of IL-6 (a), IFN-a (b), IFN-g (c), IL-8 (d), perforin (e), IP-10 (f), MIP1a (g), MIP1b (h), IL-12p40 (i), IL-18 (j), TNF-a (k) and IL-1Ra (l)are expressed in Log10 concentration in picogram per mL of plasma. (red – 0 dpi; purple – 1 dpi; blue – 2 dpi; green – 3 dpi). (n=4) Data is represented as mean± SEM. One way repeated-measures ANOVA with Geisser-Greenhouse correction for sphericity and Tukey’s post hoc correction for multiple-testing (GraphPad Prism 8) was applied. * P<0.005, ** P<0.005, *** P<0.0005.

**Figure S7.**
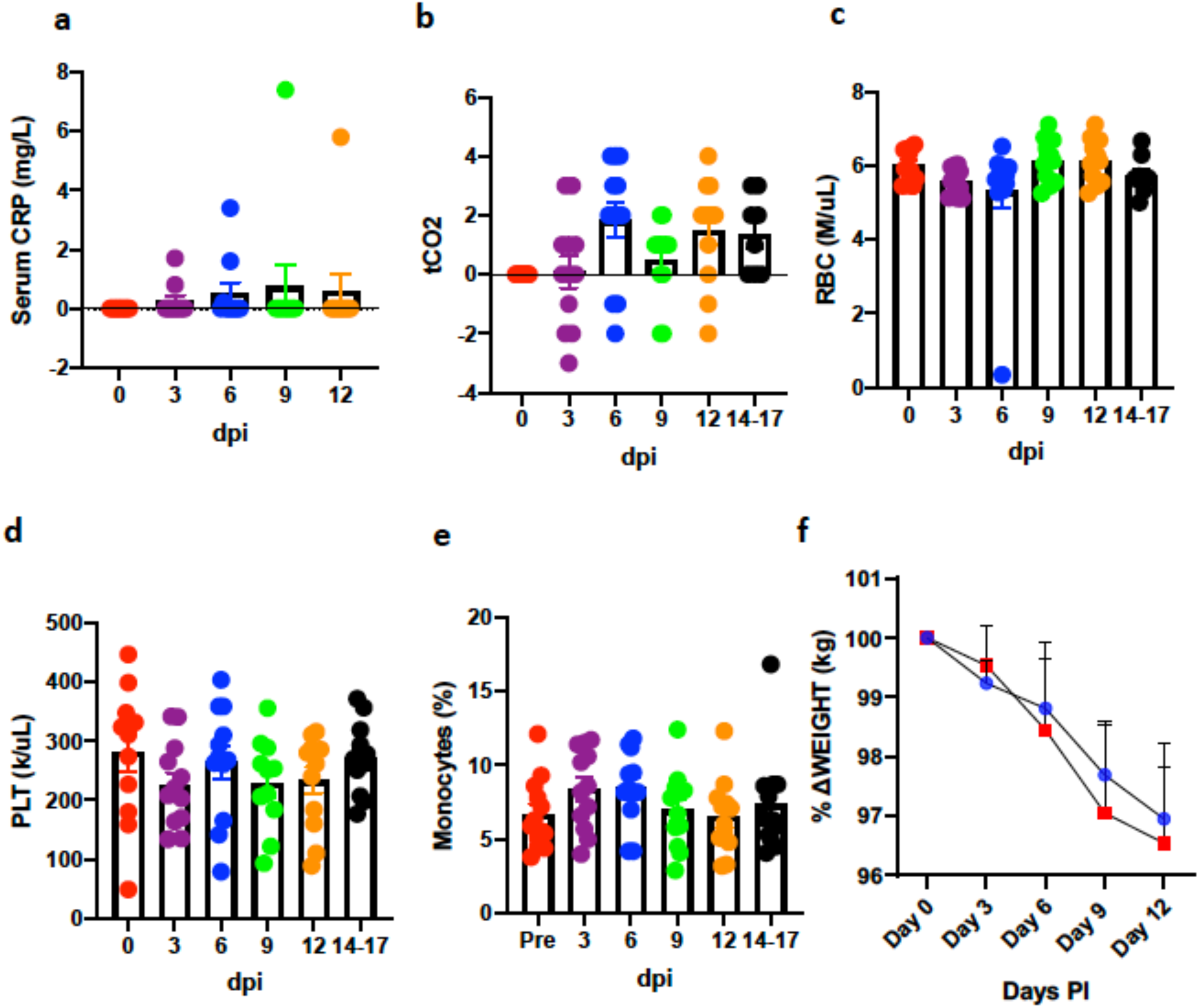
Clinical correlates in long-term (14-17 dpi) rhesus macaques. Serum levels of CRP (mg/L) (a), tCO2 (D-mmol/L) (b), and whole blood levels of Red Blood Cells (RBCs) (million/mL) (c), reticulocytes (K/mL) (d), percentage of Neutrophils (g), Neutrophils (K/mL) (f), platelets (K/uL) (e), percentage of monocytes (h) and percent change in weight (i) (Coloring scheme for I – young (blue), old (red).). (a-e) (n=12) Data is represented as mean± SEM. One way repeated-measures ANOVA with Geisser-Greenhouse correction for sphericity and Tukey’s post hoc correction for multiple-testing (GraphPad Prism 8) was applied. * P<0.005, ** P<0.005, *** P<0.0005.

**Figure S8.**
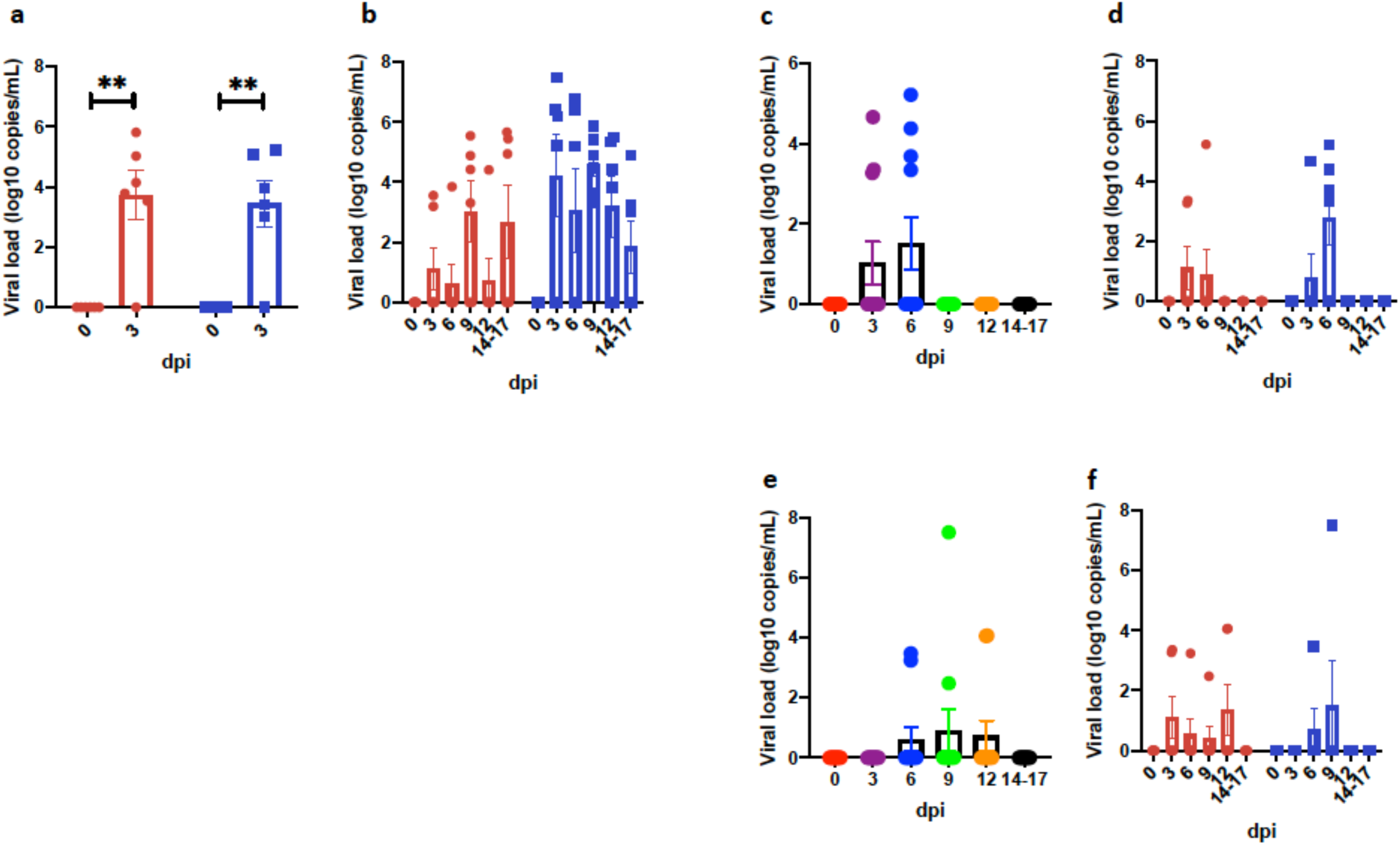
**Longitudinal viral RNA determination following SARS-CoV-2 infection in rhesus macaques**. Viral RNA (log_10_ copies/mL measured by RT-PCR in BAL fluid (a) and nasopharyngeal (b), buccopharyngeal (c-d) and rectal ^30^ swabs longitudinally. Data is depicted as combined for age (c,e) and data split by age a; b; d; f). Coloring scheme for c; e – (red – 0 dpi; purple – 3 dpi; blue – 6 dpi: green – 9 dpi; orange – 12 dpi: black – 14-17 dpi). (n=12) One way Repeated-measures ANOVA with Geisser-Greenhouse correction for sphericity and Tukey’s post hoc correction for multiple-testing (GraphPad Prism 8) was applied. * P<0.005, ** P<0.005, *** P<0.0005. Coloring scheme for a; b; d; f – young (blue), old (red). (n=6) Data is represented as mean± SEM. Two way Repeated-measures ANOVA with Geisser-Greenhouse correction for sphericity and Tukey’s post hoc correction for multiple-testing (GraphPad Prism 8) was applied. ** P<0.005.

**Figure S9.**
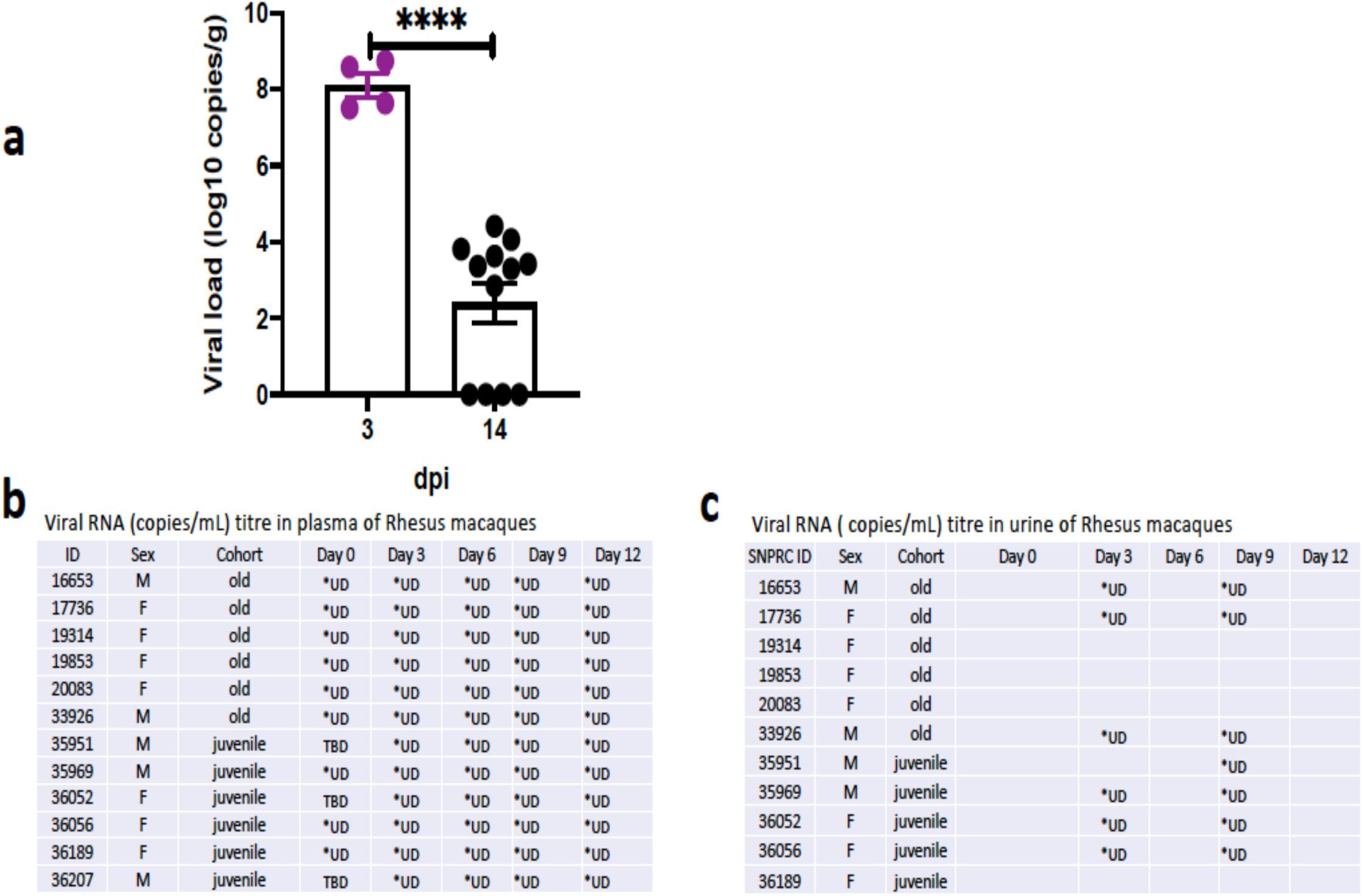
**Longitudinal viral RNA determination following SARS-CoV-2 infection in rhesus macaques**. Viral RNA was determined at endpoint in Lungs (a) and longitudinally in plasma (b) and urine (c).

**Figure S10.**
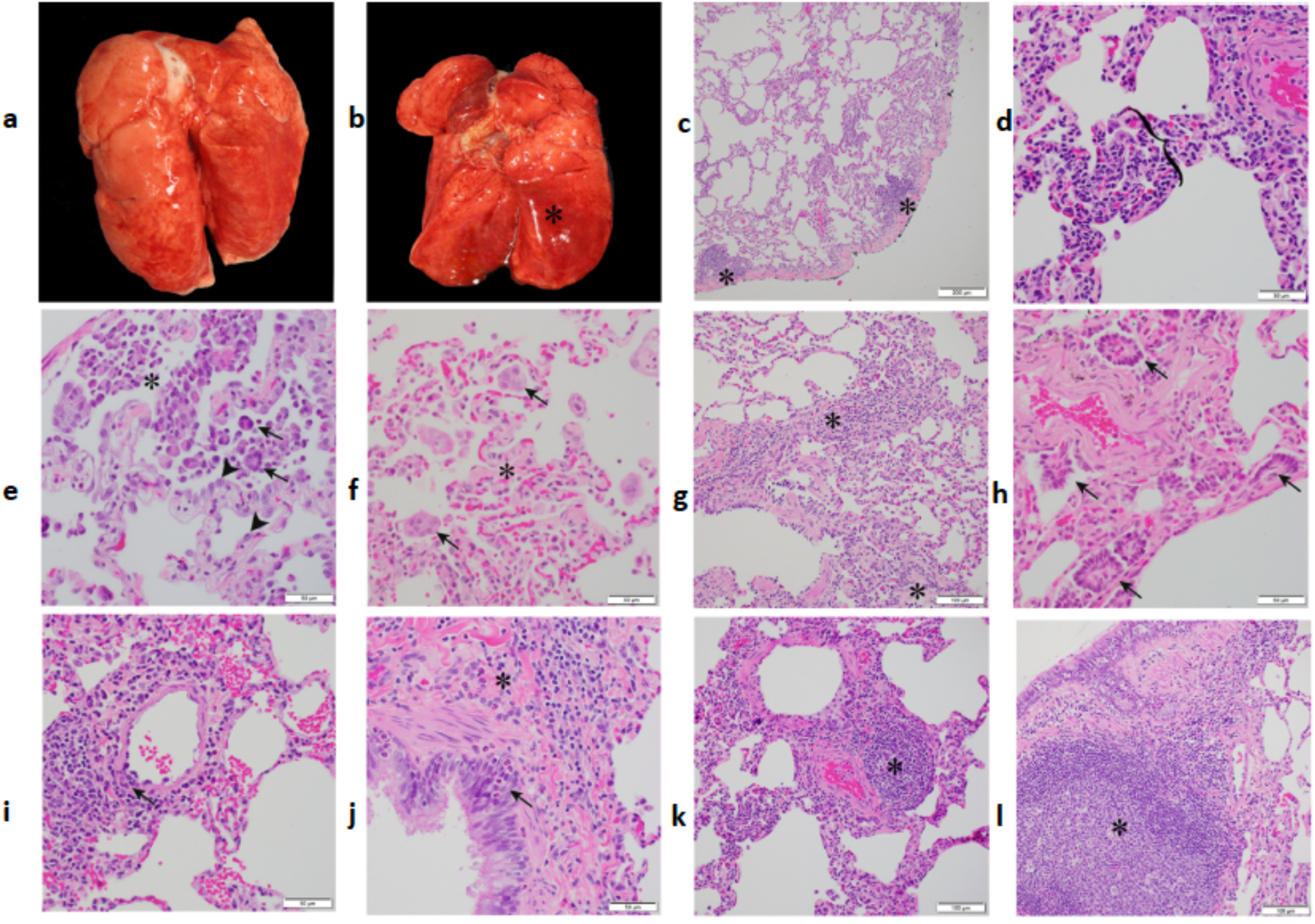
**Gross and histopathologic findings of young and aged male and female Rhesus macaques experimentally exposed to SARS-CoV-2 - 14-17 dpi**. Young male Rhesus macaque. Lung was grossly unremarkable (a). Aged male Rhesus macaque. The dorsal aspect of the lungs was mottled red (b). Young male Rhesus macaque. Lung. Subgross image showing multifocal areas of minimal interstitial pneumonia (*) (c). Young female Rhesus macaque. Lung. Mild lymphocytic interstitial pneumonia with alveolar septa (bracket) expanded by mononuclear cells (lymphocytes and macrophages) (d). Aged female Rhesus macaque. Lung. Mild lymphocytic interstitial pneumonia with increased alveolar macrophages and few syncytial cells (arrow) within the alveolar lumen (*; a neutrophil is just to the left of the *) and type II pneumocytes lining alveoli (arrowhead) (e). Aged female Rhesus macaque. Lung. Minimal interstitial pneumonia with alveolar septa expanded by fibrosis (*) and few syncytial cells (arrow) within alveoli (f). Young male Rhesus macaque. Lung. Alveolar septa expanded by fibrosis (*) and lymphocyte infiltrates (g). Aged male Rhesus macaque. Lung. Areas of bronchiolization (arrows) (h). Young female Rhesus macaque. Lung. Vasculitis. Vascular wall disrupted by infiltrates of mononuclear cells and lesser neutrophils (arrow) (i). Young female Rhesus macaque. Lung. Bronchitis. Bronchial epithelium infiltrated by eosinophils (arrow). Fibrosis adjacent to bronchus (*) (j). Young female Rhesus macaque. Lung. Area of perivascular lymphocyte infiltrates (*) (k). Young female Rhesus macaque. Lung. Area of bronchiolar associated lymphoid tissue (BALT) (*) (l). All slides were stained with H&E.

**Figure S11.**
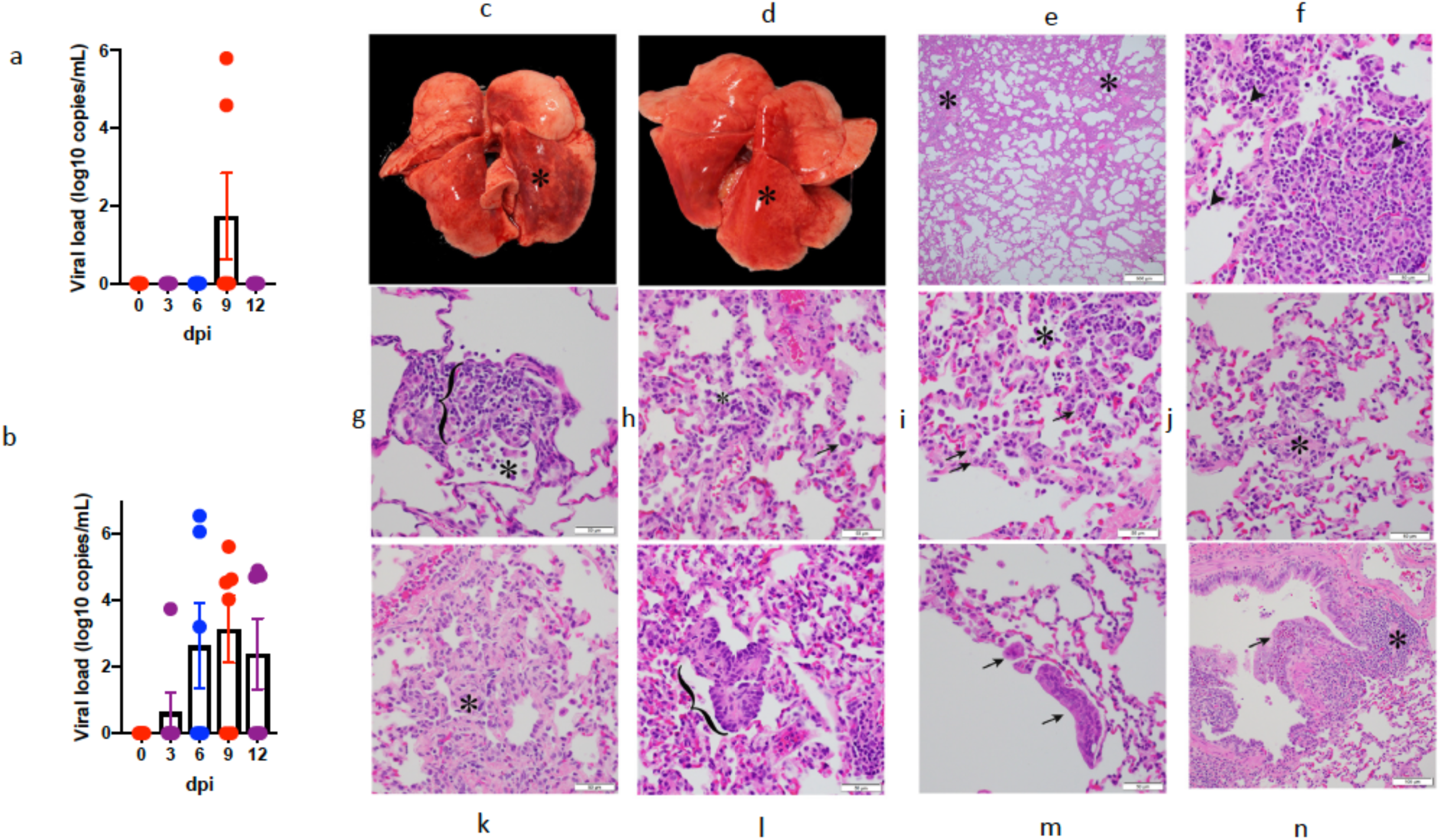
**Viral, Gross and histopathologic findings of young male and female baboons experimentally exposed to COVID19 - 14-17 dpi**. Viral RNA (log_10_ copies/mL were measured by RT-PCR in buccopharyngeal (a) and rectal (b) swabs longitudinally. Young male baboon. The dorsal aspect of the lungs was mottled red (*) (c). Young female baboon. The dorsal aspect of the lungs was mottled red (*) (d). Young male baboon. Lung. Subgross image showing areas of consolidation (*) (e). Young female baboon. Moderate lymphocytic interstitial pneumonia with scattered neutrophils (arrowhead) (f). Young female baboon. Moderate lymphocytic interstitial pneumonia with alveolar septa (bracket) markedly expanded by mononuclear cells (lymphocytes and macrophages) and increased alveolar macrophages within the alveolar lumen (*) (g). Young male baboon. Lung. Mild lymphocytic interstitial pneumonia with increased alveolar macrophages and few syncytial cells (arrow) within the alveolar lumen (*) (h). Young female baboon. Mild lymphocytic interstitial pneumonia with scattered type II pneumocytes (arrows) and increased alveolar macrophages and neutrophils within the alveolar lumen (*) (i). Young male baboon. Lung. Alveolar septa expanded by fibrosis (*) (j). Young male baboon. Lung. Alveolar septa expanded by fibrosis (*) (k). Young female baboon. Area of bronchiolization (bracket) (l). Young male baboon. Lung. Syncytial cells within airways (arrows) (m). Young male baboon. Lung. Bronchitis. Bronchial wall expanded by infiltrates of eosinophils that expand and disrupt the epithelium (arrow). Area of bronchiolar associated lymphoid tissue (BALT) (*) (n). All slides were stained with H&E.

**Figure S12.**
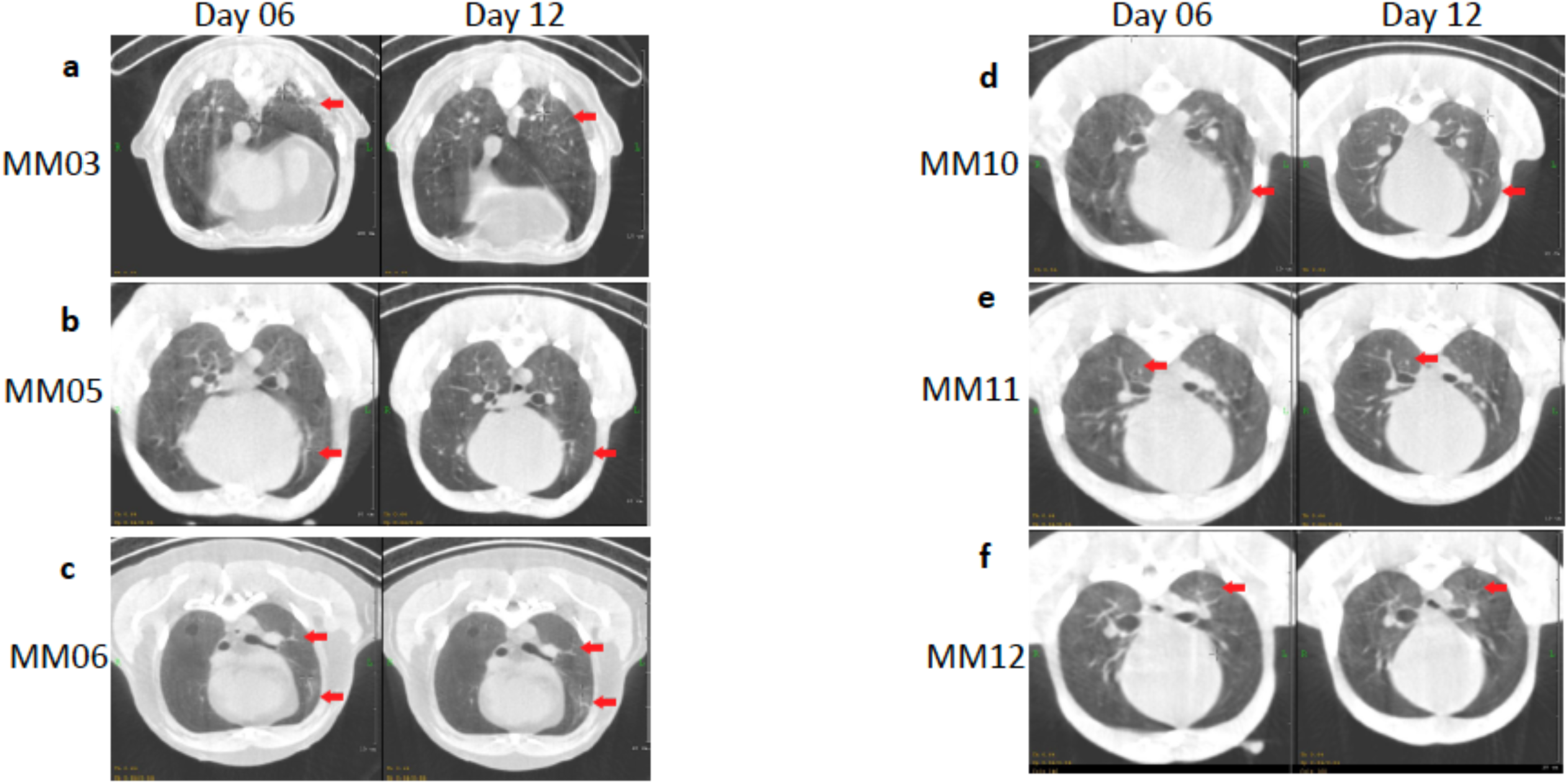
CT scan in axial view showing lesion characteristics in rhesus macaques infected with SARS-CoV-2 from Day 6-12 dpi. As seen in panel A, B, D, E and F patchy alveolar patterns, nodular and/or multifocal ground glass opacities (red arrow) seen on Day 6 dpi show dramatic resolution by Day 12 dpi, whereas panel C shows persistent patchy ground glass opacity on Day 6 dpi and Day 12 dpi.

**Figure S13.**
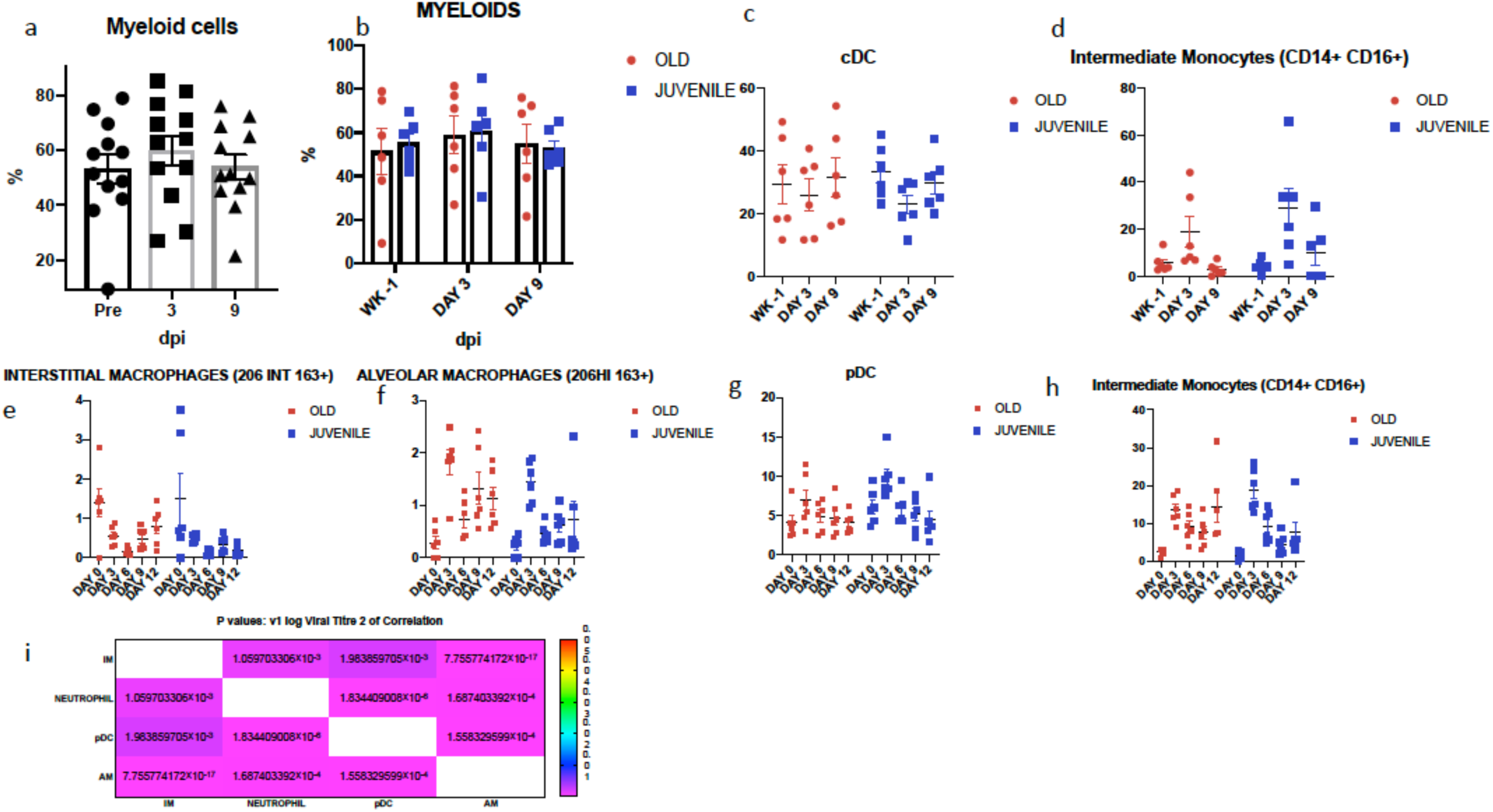
Accumulation of various types of myeloid cells in BAL (a-d) and PBMCs (c-h). Total myeloid cell compartment in the BAL in all animals (a) (n=12), and in two groups of macaques split by age (b). percentage of cDCs (c) and intermediate monocytes (d) in BAL. Percentage of interstitial (e) and alveolar (f) macrophages, pDCs (g) and intermediate macrophages (h) in the peripheral blood. Coloring scheme for b-h – young (blue), old (red) (n=6). (i) P value table for Spearman’s correlation curve in Fig 5i.

**Figure S14.**
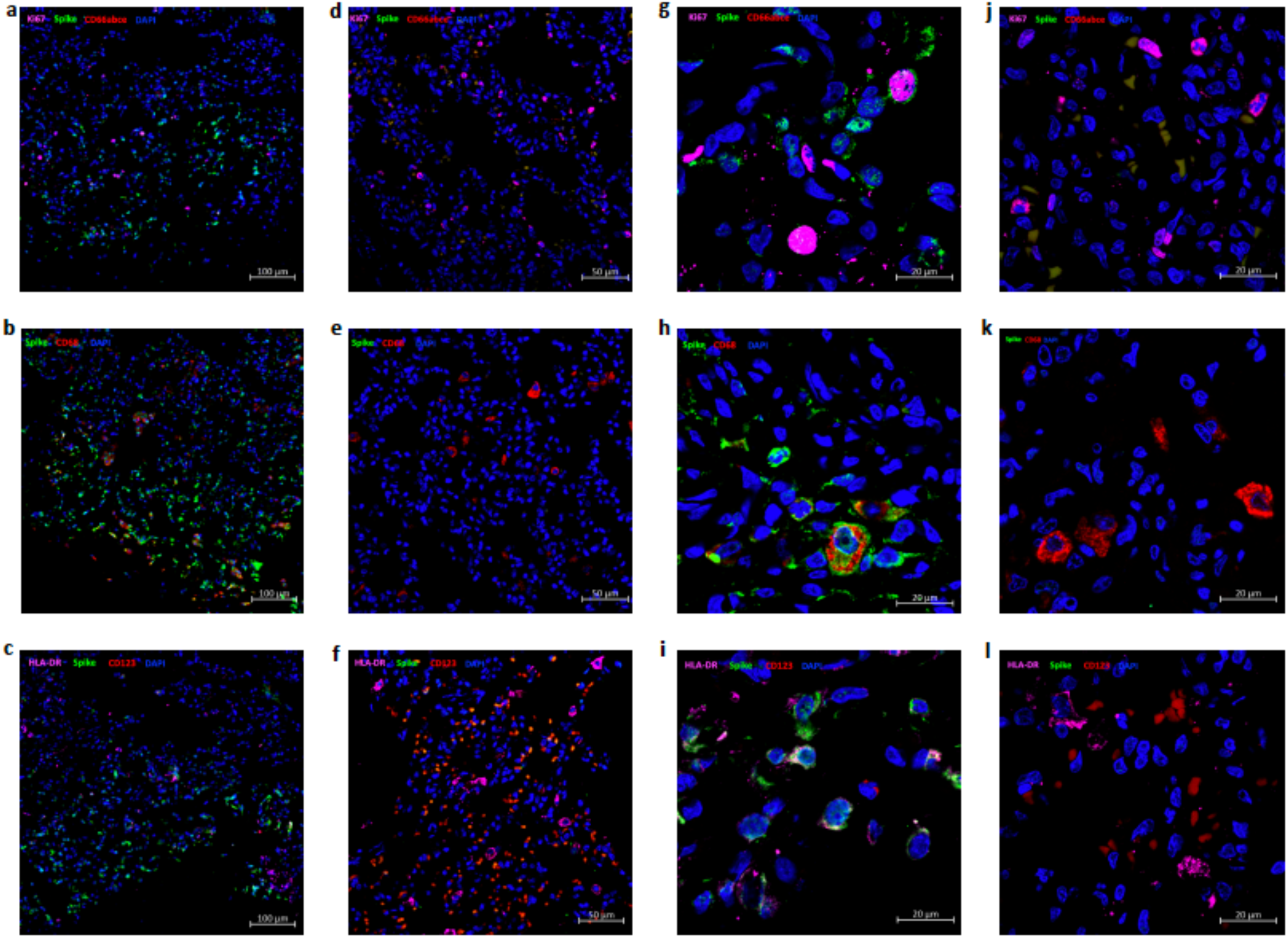
**Detection of SARS-CoV-2 signal in host lung cells by confocal microscopy**. Multi-label confocal immunofluorescence microscopy of a high viral titer lung lobe from SARS CoV-2 infected Rhesus macaque at 3 dpi with SARS CoV-2 Spike specific antibody (green), neutrophil marker CD66abce (red) and DAPI (blue)- (10X-a, 63X-g) vs the naïve control lungs (10X-d, 63X-j). SARS CoV-2 Spike (green), pan-macrophage marker CD68 (red) and DAPI (blue) in infected lungs (10X-b and 63X-h) vs the naïve control lungs (10X-e, 63X-k). SARS CoV-2 Spike (green), HLA-DR ^29^, pDC marker CD123 (red) and DAPI (blue) specific staining in infected lungs (10X-c,63X-i) vs naïve control lungs(10X-f, 63X-l).

**Figure S15.**
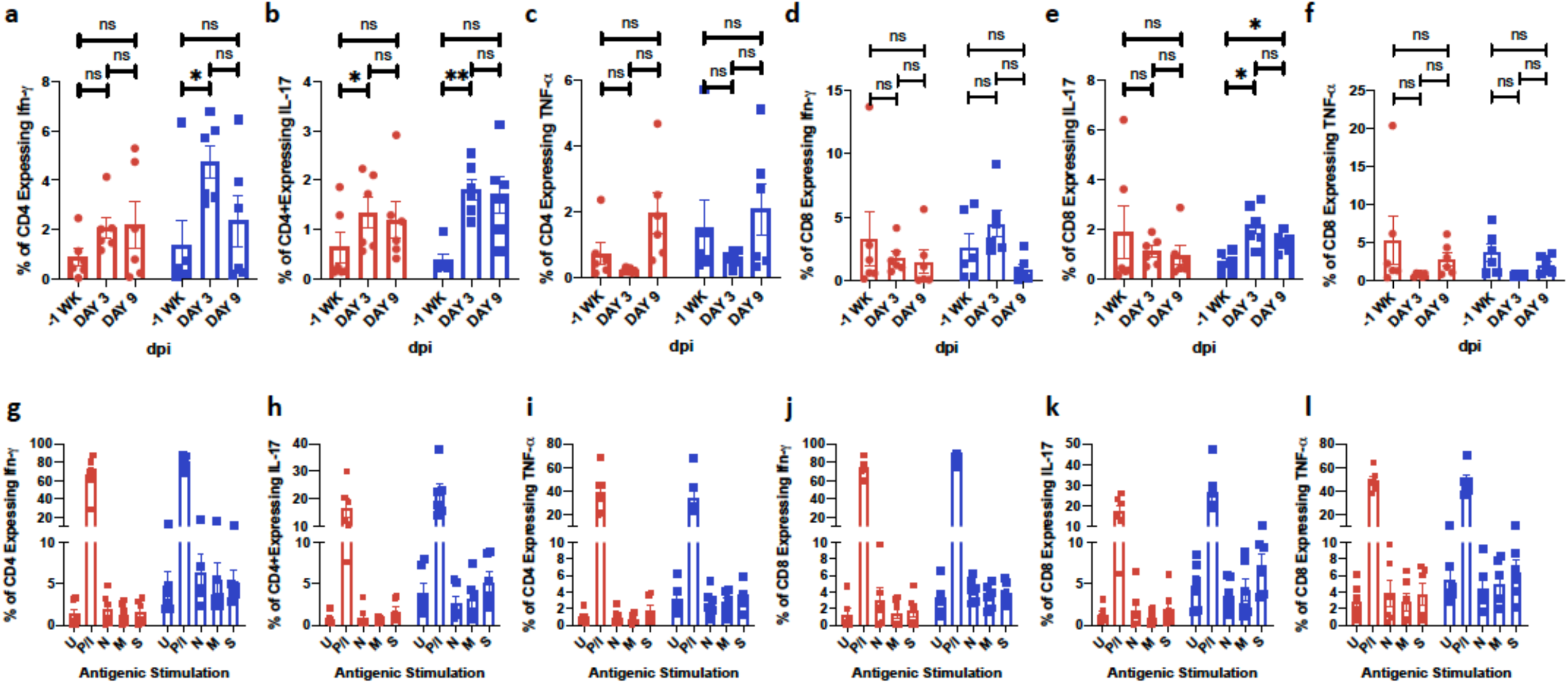
**Longitudinal changes in cytokine secretion profile in BAL T cells following SARS-CoV-2 infection in rhesus macaques**. BAL Frequencies of CD4^+^ T cell subsets expressing Interferon-γ (a), IL-17 (b), TNF-α (c), CD8+ T cells expressing Interferon-γ (d), IL-17 (e), TNF-α (f) cultured overnight without any external antigenic stimulation. BAL cells were also stimulated overnight (12-14 hours) with either Mock control (U); PMA-Ionomycin (P/I) or SARS-CoV-2 -specific peptide pools of the nucleocapsid (N), membrane (M) and spike (S) proteins. Antigen specific cytokine secretion in T cells was estimated by flow cytometry. Fraction of CD4+ T cell subsets expressing Interferon-*γ* (g), IL-17 (h), TNF-*α* (i), CD8+ T cells expressing Interferon-*γ* (j), IL-17 (k), TNF-*α* (l). Coloring scheme– young (blue), old (red). Data is represented as mean± SEM. (n=6) Two way Repeated-measures ANOVA with Geisser-Greenhouse correction for sphericity and Tukey’s post hoc correction for multiple-testing (GraphPad Prism 8) was applied. * P<0.005, ** P<0.005, *** P<0.0005.

**Figure S16.**
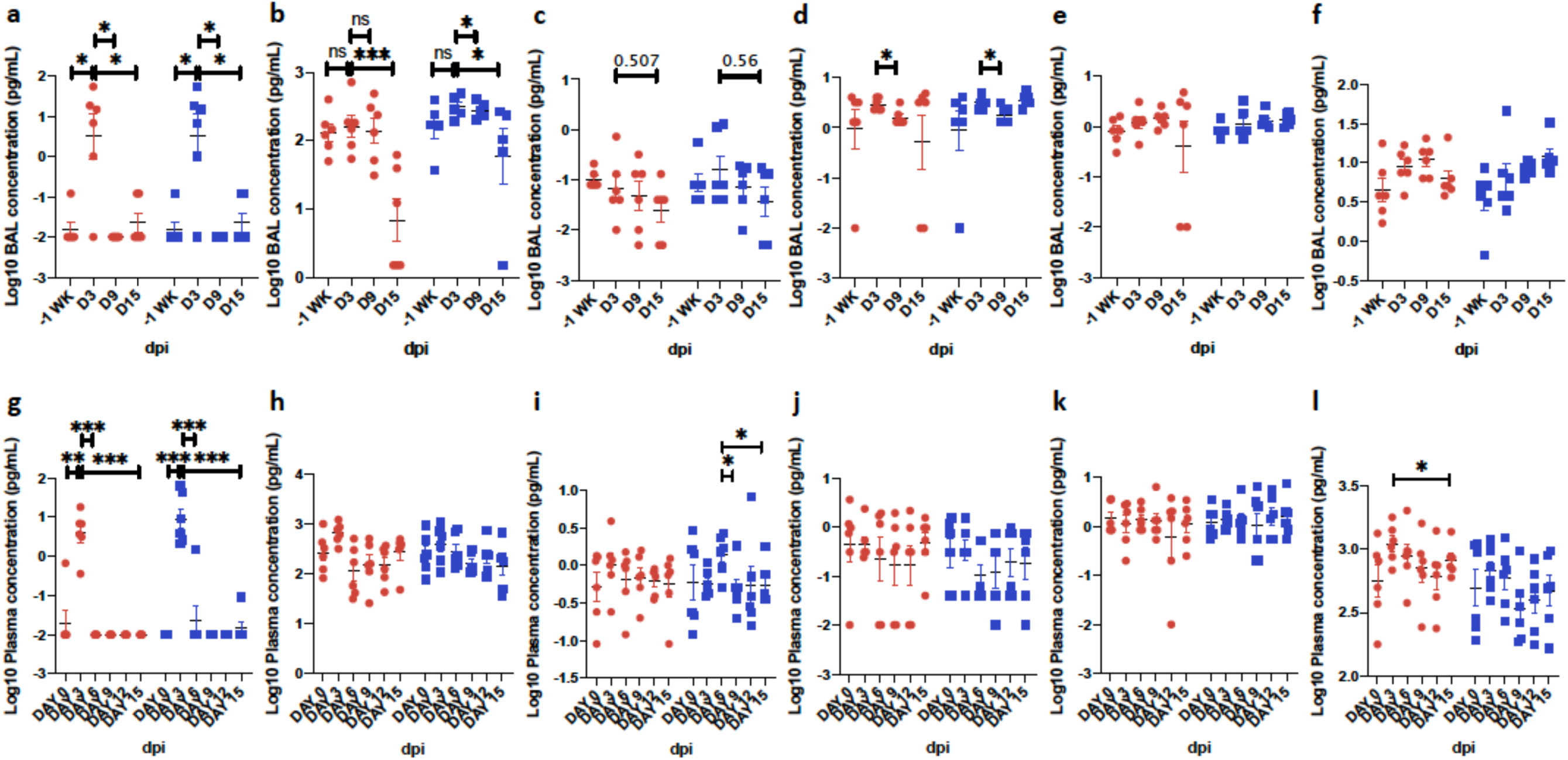
**Longitudinal changes in SARS-CoV-2 induced cytokines in BAL fluid and plasma following SARS-CoV-2 infection in rhesus macaques over two weeks**. Simultaneous analysis of multiple cytokines by Luminex technology in the BAL fluid and plasma of rhesus macaques over 0-15 dpi. Levels of IFN-*α* (a), IL-1Ra (b), IFN-*γ* (c), TNF-*α* (d), IL-6 (e), Perforin (f) are expressed in Log10 concentration in picogram per mL of BAL fluid. Levels of IFN-*α* (g), IL-1Ra (h), IFN-*γ* (i), TNF-a (j), IL-6 (k), Perforin (l) are expressed in Log10 concentration in picogram per mL of BAL fluid. Coloring scheme – young (blue), old (red). Data is represented as mean± SEM. (n=12) Two way Repeated-measures ANOVA with Geisser-Greenhouse correction for sphericity and Tukey’s post hoc correction for multiple-testing (GraphPad Prism 8) was applied. * P<0.005, ** P<0.005, *** P<0.0005.

**Figure S17.**
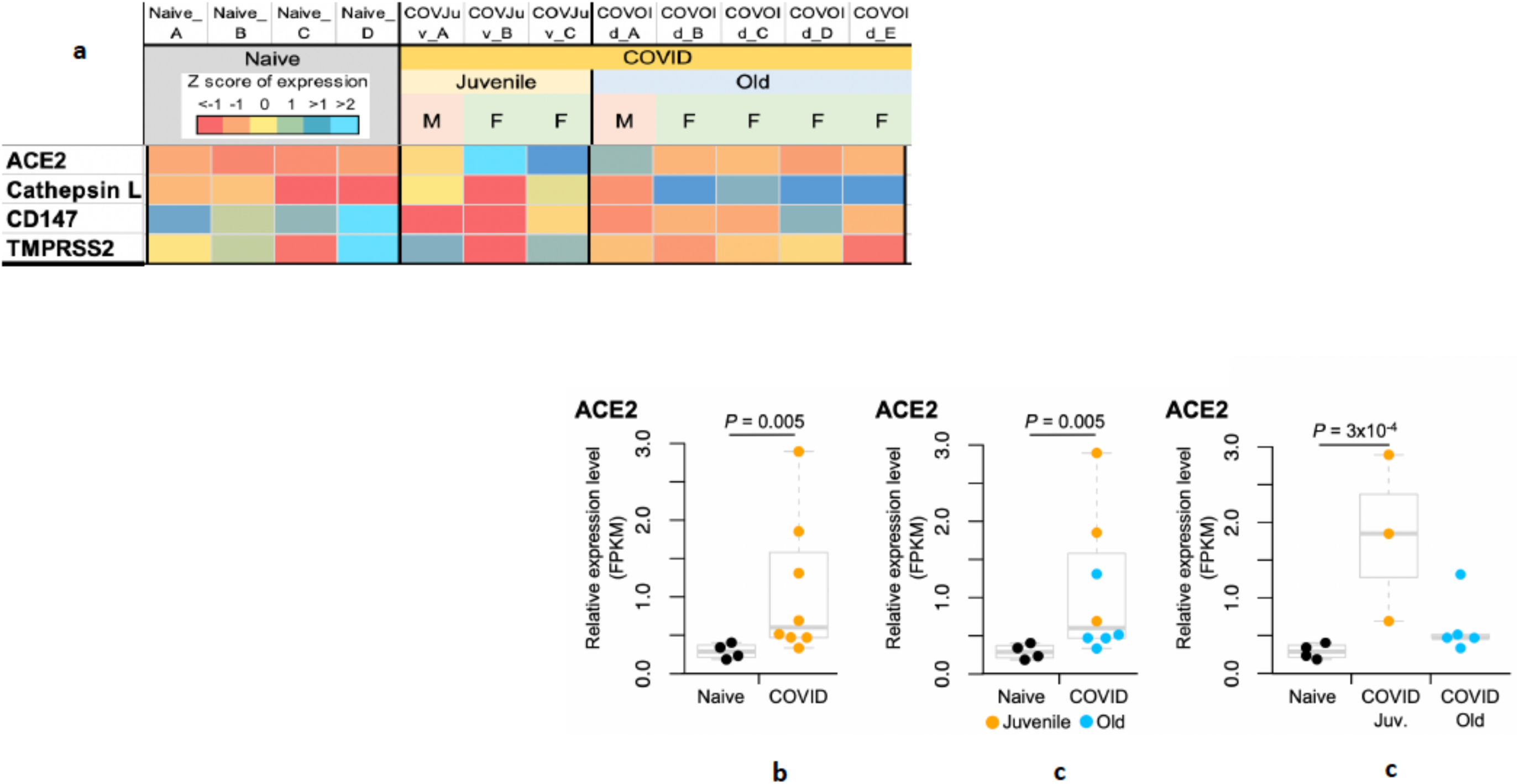
**SARS-CoV-2 infection induces ACE-2 expression**. RNAseq was performed on total RNA isolated from the lungs of naïve (n=3) and SARS-CoV-2 infected (14-17dpi) rhesus macaques (n=8, 3 young and 5 old macaques) as described earlier ^16^. Results indicate that the expression of ACE2, which is lower in naïve animals (denoted by red color in the heat map) (a), was induced following SARS-CoV-2 infection (denoted by blue color in the heat map) (a). Relative expression level of ACE-2 was significantly higher than in naïve tissues (b). Higher expression of ACE-2 was observed in lung tissues obtained at necropsy from young relative to old macaques (c, d), such that the difference between naïve animals and young SARS-CoV-2 infected animals in ACE-2 expression levels was statistically significant by itself. All p-values shown on expression swarm plots (b-d) are FDR-corrected significance values for differential expression calculated by DESEQ2^16^.

**Figure S18.**
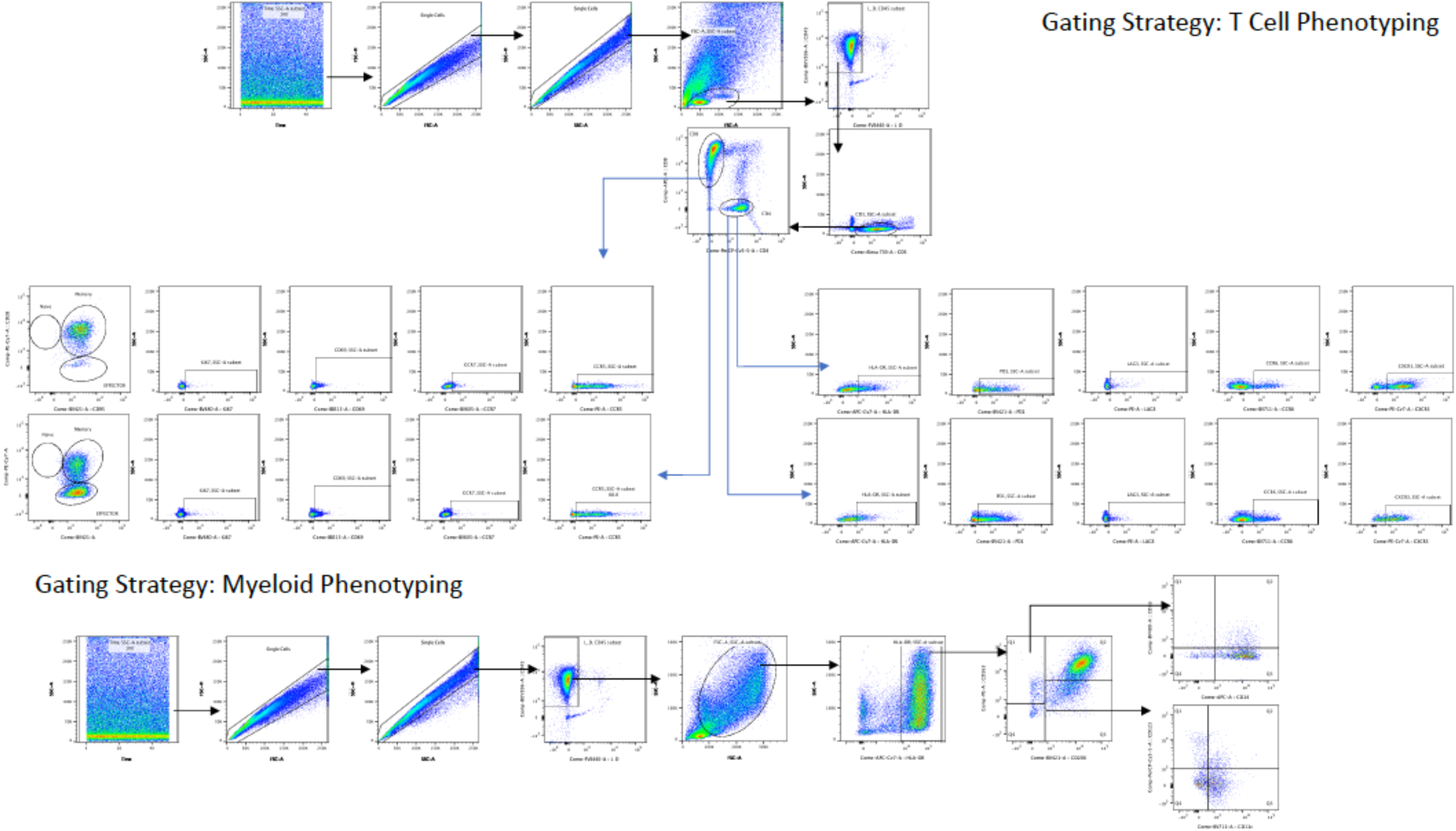
**Flow cytometry Gating Strategy**. Gating strategy for T cell phenotyping is described.

## Supplemental tables

Table S1. In vivo experimental design. A. short-term rhesus macaque pilot. B-D. 14-day multispecies comparison in rhesus macaques, baboons, marmosets.

Table S2. Distribution of lesions by anatomic location and morphologic diagnosis of young and aged rhesus macaques experimentally exposed to SARS-CoV-2 - 3 dpi.

Table S3. Distribution of lesions by anatomic location and morphologic diagnosis of young and aged rhesus macaques experimentally exposed to SARS-CoV-2 – 14-17 dpi.

Table S4. Distribution of lesions in baboons experimentally exposed to SARS-CoV-2 – 14-17 dpi.

Table S5. CXR scores in rhesus macaques experimentally exposed to SARS-CoV-2 – 14-17 dpi.

Table S6. CT scores in rhesus macaques experimentally exposed to SARS-CoV-2 – 14-17 dpi.

Table S7. List of antibodies used for immunophenotyping studies.

