## Supplemental Tables S1-S7 for "SARS-CoV-2 infection leads to acute infection with dynamic cellular and inflammatory flux in the lung that varies across nonhuman primate species"

| Table S1. In vivo experimental design. A. short-term rhesus macaque pilot. B-D. 14-day multispecies comparison in rhesus macaques, baboons, marmosets. | | | | | | |
|---|---|---|---|---|---|---|
| **Id** | **Species** | **Common name** | **Gender** | **Age (Year and Days)** | **Endpoint We** | **MHC Summary** |
| MM01 | Macaca mulatta | Rhesus monkey | Male | 22 years, 103 days | 9.9 | A001: NEGATIVE B003: NEGATIVE B008: NEGATIVE B017: NEGATIVE |
| MM02 | Macaca mulatta | Rhesus monkey | Male | 21 years, 181 days | 10.26 | A001: NEGATIVE B003: NEGATIVE B008: NEGATIVE B017: NEGATIVE |
| MM03 | Macaca mulatta | Rhesus monkey | Female | 17 years, 335 days | 7.6 | A001: POSITIVE B003: NEGATIVE B008: NEGATIVE B017: NEGATIVE |
| MM04 | Macaca mulatta | Rhesus monkey | Female | 19 years, 127 days | 8.5 | A001: NEGATIVE B003: NEGATIVE B008: NEGATIVE B017: POSITIVE |
| MM05 | Macaca mulatta | Rhesus monkey | Female | 16 years, 148 days | 7.06 | A001: POSITIVE B003: NEGATIVE B008: POSITIVE B017: NEGATIVE |
| MM06 | Macaca mulatta | Rhesus monkey | Female | 15 years, 299 days | 8.98 | A001: NEGATIVE B003: NEGATIVE B008: NEGATIVE B017: NEGATIVE |
| MM07 | Macaca mulatta | Rhesus monkey | Female | 15 years, 124 days | 5.5 | A001: NEGATIVE B003: NEGATIVE B008: NEGATIVE B017: NEGATIVE |
| MM08 | Macaca mulatta | Rhesus monkey | Male | 17 years, 349 days | 8.16 | A001: NEGATIVE B003: NEGATIVE B008: NEGATIVE B017: NEGATIVE |
| MM09 | Macaca mulatta | Rhesus monkey | Male | 3 years, 350 days | 6. | A001: POSITIVE B003: NEGATIVE B008: POSITIVE B017: NEGATIVE |
| MM10 | Macaca mulatta | Rhesus monkey | Male | 3 years, 343 days | 6.5 | A001: POSITIVE B003: NEGATIVE B008: NEGATIVE B017: NEGATIVE |
| MM11 | Macaca mulatta | Rhesus monkey | Female | 3 years, 323 days | 5.84 | A001: POSITIVE B003: NEGATIVE B008: NEGATIVE B017: NEGATIVE |
| MM12 | Macaca mulatta | Rhesus monkey | Female | 3 years, 324 days | 4.44 | A001: NEGATIVE B003: NEGATIVE B008: NEGATIVE B017: POSITIVE |
| MM13 | Macaca mulatta | Rhesus monkey | Female | 3 years, 316 days | 4.21 | A001: NEGATIVE B003: NEGATIVE B008: NEGATIVE B017: POSITIVE |
| MM14 | Macaca mulatta | Rhesus monkey | Female | 3 years, 269 days | 5.7 | A001: POSITIVE B003: NEGATIVE B008: NEGATIVE B017: NEGATIVE |
| MM15 | Macaca mulatta | Rhesus monkey | Male | 3 years, 258 days | 5.72 | A001: NEGATIVE B003: NEGATIVE B008: NEGATIVE B017: POSITIVE |
| MM16 | Macaca mulatta | Rhesus monkey | Male | 3 years, 19 days | 4.81 | A001: NEGATIVE B003: NEGATIVE B008: NEGATIVE B017: POSITIVE |
| CS01 | Callithrix spp. | Marmoset | Male | 11 years, 240 days | 0.35 | |
| CS02 | Callithrix spp. | Marmoset | Male | 6 years, 292 days | 0.41 | |
| CS03 | Callithrix spp. | Marmoset | Female | 7 years, 22 days | 0.42 | |
| CS04 | Callithrix spp. | Marmoset | Female | 6 years, 235 days | 0.36 | |
| CS05 | Callithrix spp. | Marmoset | Male | 6 years, 220 days | 0.35 | |
| CS06 | Callithrix spp. | Marmoset | Female | 6 years, 100 days | 0.51 | |
| PH01 | Papio hamadryas anubis/Papio hamadryas cynocephalu | Olive/Yellow baboon | Female | 2 years, 104 days | 7.16 | |
| PH02 | Papio hamadryas anubis/Papio hamadryas cynocephalu | Olive/Yellow baboon | Male | 2 years, 98 days | 8.2 | |
| PH03 | Papio hamadryas anubis | Olive baboon | Female | 2 years, 76 days | 6.92 | |
| PH04 | Papio hamadryas anubis | Olive baboon | Male | 2 years, 58 days | 6.94 | |
| PH05 | Papio hamadryas anubis | Olive baboon | Female | 2 years, 36 days | 5.6 | |
| PH06 | Papio hamadryas anubis/Papio hamadryas cynocephalu | Olive/Yellow baboon | Male | 2 years, 17 days | 7.26 | |

| Table S2. Distribution of lesions by anatomic location and morphologic diagnosis of Young and Aged (Male and Female) Rhesus Macaques Experimentally Exposed to COVID19 - 3 days PE | | | | |
|---|---|---|---|---|
| **Tissue** | **Morphologic Diagnosis/es** | **Total Lesions** | **Young Total (M/F) n=2** | **Aged Total (M/F) n=2** |
| Lung | Anthracosis | 4 | 1/1 | 1/1 |
| | Interstitial mononuclear inflammation | 4 | 1/1 | 1/1 |
| | Increased alveolar histiocytes | 4 | 1/1 | 1/1 |
| | Alveolar syncytia | 4 | 1/1 | 1/1 |
| | Bronchitis | 4 | 1/1 | 1/1 |
| | BALT hyperplasia | 4 | 1/1 | 1/1 |
| | Type II pneumocyte hyperplasia | 4 | 1/1 | 1/1 |
| | Interstitial or Intralveolar neutrophilic and eosinophilic inflammation | 4 | 1/1 | 1/1 |
| | Fibrosis | 4 | 1/1 | 1/1 |
| | Vasculitis | 3 | 0/1 | 1/1 |
| | Fibrin | 3 | 1/0 | 1/1 |
| | Bronchiolitis | 2 | 1/1 | 0/0 |
| | Bronchiolization | 2 | 0/0 | 1/1 |
| | Consolidation | 2 | 1/1 | 0/0 |
| | Edema | 2 | 1/0 | 0/1 |
| | Antherosclerosis | 1 | 0/0 | 1/0 |
| Liver | Lipogranulomas | 3 | 0/1 | 1/1 |
| | Hepatitis | 2 | 1/0 | 1/0 |
| | Vacuolation, glycogen | 2 | 1/0 | 0/1 |
| | Vacuolation, lipid | 1 | 0/0 | 0/1 |
| Heart | Myocarditis | 3 | 0/1 | 1/1 |
| | Cardiomyopathy | 2 | 0/0 | 1/1 |
| | Atherosclerosis | 1 | 0/0 | 0/1 |
| Lymph Node, Mandibular | Hyperplasia | 2 | 1/0 | 1/0 |
| | Erythrophagocytosis | 2 | 1/0 | 1/0 |
| | Lymphadenitis | 1 | 0/1 | 0/0 |

| | | | | |
|---|---|---|---|---|
| Lymph Node, Inguinal | Hyperplasia | 2 | 1/1 | 0/0 |
| | Peri-nodal inflammation | 1 | 0/0 | 0/1 |
| | Erythrophagocytosis | 1 | 0/0 | 0/1 |
| Trachea | Tracheitis | 4 | 1/1 | 1/1 |
| Spleen | Hyperplasia | 3 | 1/1 | 1/0 |
| | Neutrophils red pulp | 1 | 0/1 | 0/0 |
| Stomach | Gastritis | 3 | 1/1 | 1/0 |
| | Atherosclerosis | 1 | 0/0 | 0/1 |
| Rectum | Vasculitis | 1 | 0/0 | 1/0 |
| | Myositis | 1 | 0/0 | 1/0 |
| | Myenteric plexus inflammation | 1 | 0/0 | 1/0 |
| | GALT hyperplasia | 1 | 0/1 | 0/0 |
| Nasal cavity | Rhinitis | 4 | 1/1 | 1/1 |
| Kidney | Glomerulopathy | 2 | 0/0 | 1/1 |
| | Atherosclerosis | 1 | 0/0 | 1/0 |
| Duodenum | Serositis | 1 | 0/0 | 1/0 |
| | GALT hyperplasia | 1 | 0/0 | 0/1 |
| | Myenteric plexus inflammation | 1 | 0/1 | 0/0 |
| Jejunum | Myositis | 1 | 0/0 | 1/0 |
| | Myenteric plexus inflammation | 1 | 0/0 | 1/0 |
| | GALT hyperplasia | 1 | 0/0 | 1/0 |
| Eye | Choroiditis | 1 | 0/0 | 1/0 |
| | Conjunctivitis | 1 | 0/1 | 0/0 |
| | Descemet's membrane duplication with abnormal corneal endothelium | 1 | 0/0 | 0/1 |
| Lymph Node, Axillary | Hyperplasia | 2 | 0/1 | 1/0 |
| Lymph node, mediastinal | Hyperplasia | 1 | 1/0 | 0/0 |

| | | | | |
|---|---|---|---|---|
| | Erythrophagocytosis | 1 | 1/0 | 0/0 |
| Adrenal Gland | Amyloid | 1 | 0/0 | 0/1 |
| | Mineralization | 1 | 0/0 | 0/1 |
| Colon | Colitis | 2 | 0/0 | 1/1 |
| Salivary Gland, Mandibular | Sialadenitis | 1 | 1/0 | 0/0 |
| Tonsil | Tonsillitis | 1 | 0/0 | 1/0 |
| Ileum | GALT hyperplasia | 1 | 1/0 | 0/0 |
| Brain | Vacuolation, white matter | 1 | 0/0 | 1/0 |
| This table will exclude tissues that were normal in all young and old animals (skin, thyroid gland, ovary, testes) | | | | |

| Table S3. Distribution of lesions by anatomic location and morphologic diagnosis of Young and Aged (Male and Female) Rhesus Macaques Experimentally Exposed to COVID19 - 2 weeks PE | | | | |
|---|---|---|---|---|
| Tissue | Morphologic Diagnose/es | Total Lesions | Young Total (M/F) | Aged Total (M/F) |
| Lung | Anthracosis | 12 | 6(3/3) | 6(2/4) |
| | Syncytia | 12 | 6(3/3) | 6(2/4) |
| | Interstitial mononuclear inflammation | 11 | 6(3/3) | 5(1/4) |
| | Increased alveolar histiocytes | 9 | 5(3/2) | 4(0/4) |
| | Prominent perivascular lymphocytes | 7 | 5(2/3) | 2(0/2) |
| | Interstitial or Intralveolar neutrophilic inflammation | 5 | 3(1/2) | 2(0/2) |
| | Alveolar interstitial fibrosis | 5 | 2(1/1) | 3(0/3) |
| | BALT hyperplasia | 5 | 3(1/2) | 2(0/2) |
| | Bronchitis | 4 | 2(0/2) | 2(0/2) |
| | Type II pneumocyte hyperplasia | 4 | 2(2/0) | 2(0/2) |
| | Vasculitis | 3 | 2(1/1) | 1(1/0) |
| | Bronchiolization | 2 | 0 | 2(1/1) |
| | Bronchiolitis | 1 | 0 | 1(0/1) |
| Spleen | Lymphoid hyperplasia | 12 | 6(3/3) | 6(2/4) |
| | Neutrophilic Splenitis | 6 | 4(2/2) | 2(1/1) |
| Liver | Lipogranulomas | 8 | 4(3/1) | 4(1/3) |
| | Hepatitis | 3 | 1(0/1) | 2(1/1) |
| | Vacuolation, glycogen | 3 | 1(1/0) | 2(0/2) |
| | Perivasculitis | 1 | 1(0/1) | 0 |
| Trachea | Tracheitis | 11 | 6(3/3) | 5(2/3) |
| Lymph Node, Mandibular | Lymphoid hyperplasia | 8 | 4(3/1) | 4(2/2) |
| | Erythrophagocytosis | 1 | 0 | 1(0/1) |
| | Vasculitis | 1 | 1(0/1) | 0 |
| Lymph node, mediastinal | Lymphoid hyperplasia | 6 | 4(3/1) | 2(1/1) |
| | Anthracosis | 3 | 0 | 3(1/2) |
| | Erythrophagocytosis | 1 | 1(1/0) | 0 |

| | | | | |
|---|---|---|---|---|
| Heart | Cardiomyopathy | 3 | 0 | 3(1/2) |
| | Myocarditis | 3 | 0 | 3(1/2) |
| | Fibrosis | 2 | 0 | 2(1/1) |
| Stomach | Gastritis | 7 | 6(3/3) | 1(0/1) |
| Lymph Node, Inguinal | Lymphoid hyperplasia | 6 | 3(2/1) | 3(2/1) |
| Adrenal Gland | Mineralization | 4 | 3(1/2) | 1(1/0) |
| | Adrenalitis | 1 | 0 | 1(0/1) |
| | Vasculitis | 1 | 0 | 1(0/1) |
| Kidney | Nephritis | 2 | 1(1/0) | 1(0/1) |
| | Fibrosis | 1 | 0 | 1(1/0) |
| | Tubular ectasia | 1 | 0(0/0) | 1(1/0) |
| Lymph Node, Axillary | Lymphoid hyperplasia | 5 | 4(2/2) | 1(1/0) |
| Salivary Gland, Mandibular | Sialadenitis | 4 | 2(2/0) | 2(0/2) |
| | Adenoma | 1 | 0 | 1(0/1) |
| Thyroid | Cysts | 2 | 1(1/0) | 1(0/1) |
| | Ectopic thymus | 1 | 1(0/1) | 0 |
| | Hyperplasia | 1 | 0 | 1(0/1) |
| | Thyroiditis | 1 | 0 | 1(0/1) |
| Tonsil | Tonsillitis | 4 | 3(3/0) | 1(0/1) |
| Rectum | Proctitis | 2 | 1(0/1) | 1(1/0) |
| | Myenteric plexus inflammation | 1 | 0 | 1(1/0) |
| Colon | Colitis | 2 | 1(0/1) | 1(1/0) |
| Eye | Choroiditis | 1 | 1(0/1) | 0 |
| | Myositis, extraocular muscle | 1 | 0 | 1(1/0) |
| Nasal cavity | Rhinitis | 2 | 2(1/1) | 0 |
| Testes | Vasculitis | 1 | 0 | 1(1/na) |

This table excludes tissues that were normal in young and old animals (skin, duodenum, jejunum, ileum, ovary and brain) and  parasitic disease.

| Table S4. Distribution of lesions by anatomic location and morphologic diagnosis of Young (Male and Female) Baboons Experimentally Exposed to COVID19 - 2 weeks PE | | | | |
|---|---|---|---|---|
| Tissue | Morphologic Diagnosis/es | Total Lesions | Young Total Males n=3 | Young Total Female n=3 |
| Lung | Anthracosis | 6 | 3 | 3 |
| | Interstitial mononuclear inflammation | 6 | 3 | 3 |
| | Increased alveolar histiocytes | 6 | 3 | 3 |
| | Alveolar syncytia | 6 | 3 | 3 |
| | Bronchitis, eosinophilic | 6 | 3 | 3 |
| | BALT hyperplasia | 5 | 3 | 2 |
| | Type II pneumocyte hyperplasia | 4 | 2 | 2 |
| | Interstitial or Intralveolar neutrophilic inflammation | 3 | 2 | 1 |
| | Alveolar interstitial fibrosis | 2 | 1 | 1 |
| | Bronchiolization | 1 | 0 | 1 |
| Spleen | Increased neutrophils | 6 | 3 | 3 |
| | Lymphoid hyperplasia | 6 | 3 | 3 |
| Liver | Vacuolation, glycogen | 5 | 3 | 2 |
| | Lipogranulomas | 4 | 2 | 2 |
| | Hepatitis | 2 | 1 | 1 |
| Colon | GALT hyperplasia | 4 | 2 | 2 |
| | Eosinophilic granulomas | 2 | 0 | 2 |
| | Colitis | 2 | 0 | 2 |
| | Myenteric plexus inflammation | 1 | 0 | 1 |
| Eye | Choroiditis | 4 | 2 | 2 |
| | Descemet's membrane duplication with abnormal corneal endothelium | 2 | 1 | 1 |
| | Episcleritis | 1 | 0 | 1 |
| | Iris vascular sclerosis | 1 | 1 | 0 |
| Lymph Node, Mandibular | Lymphoid hyperplasia | 5 | 2 | 3 |
| | Capsular fibrosis | 1 | 1 | 0 |
| | Erythrophagocytosis | 1 | 1 | 0 |
| Lymph Node, Inguinal | Lymphoid hyperplasia | 6 | 3 | 3 |
| Lymph Node, Axillary | Lymphoid hyperplasia | 6 | 3 | 3 |
| Stomach | Gastritis | 5 | 3 | 2 |
| | Vasculitis, eosinophilic | 1 | 1 | 0 |
| Thyroid | Ectopic thymus | 4 | 2 | 2 |
| | Cyst | 1 | 0 | 1 |
| Lymph node, mediastinal | Lymphoid hyperplasia | 5 | 2 | 3 |

| | | | | |
|---|---|---|---|---|
| **Trachea** | Tracheitis | 5 | 2 | 3 |
| **Rectum** | GALT hyperplasia | 4 | 2 | 2 |
| | Myenteric plexus inflammation | 1 | 1 | 0 |
| **Heart** | Myocarditis | 4 | 2 | 2 |
| **Nasal cavity** | Rhinitis | 4 | 3 | 1 |
| **Duodenum** | GALT hyperplasia | 4 | 3 | 1 |
| **Tonsil** | Tonsillitis | 3 | 1 | 2 |
| | Lymphoid hyperplasia | 1 | 1 | 0 |
| **Salivary Gland, Mandibular** | Sialadenitis | 3 | 2 | 1 |
| **Adrenal Gland** | Inflammation, medulla | 2 | 1 | 1 |
| **Brain** | Edema | 2 | 1 | 1 |
| **Ileum** | Peyer's patch hyperplasia | 1 | 1 | 0 |
| **Jejunum** | Myenteric plexus inflammation | 1 | 0 | 1 |
| **Kidney** | Nephritis | 1 | 1 | 0 |
| This table excludes tissues that were normal in both young males and females animals (skin, ovary and testes). | | | | |

| Table S5. Radiograph Scoring Macaques, Two Week Study | | | | | | | | | | | | |
|---|---|---|---|---|---|---|---|---|---|---|---|---|
| Animal | Baseline[1] | | Day 3 | | Day 6 | | Day 9 | | Day 11-12 | | End of project[2] | |
| | Descriptive Score | **(Total)** | Descriptive Score | **(Total)** | Descriptive Score | **(Total)** | Descriptive Score | **(Total)** | Descriptive Score | **(Total)** | Descriptive Score | **(Total)** |
| MM09 | *0 .5RVB | **(0.5)** | *0.5RLVD | **(1)** | 0 | **(0)** | *1RVB | **(1)** | *0.5RVD | **(0.5)** | 0 | **(0)** |
| MM10 | 0 | **(0)** | *1RUB | **(1)** | *1LCD | **(1)** | 0 | **(0)** | 0 | **(0)** | 0 | **(0)** |
| MM11 | 0 | **(0)** | *0.5LUVD | **(1)** | 0 | **(0)** | 0 | **(0)** | 0 | **(0)** | 0 | **(0)** |
| MM12 | 0 | **(0)** | 1RLUVDE | **(4)** | 1RLUVD | **(4)** | 1RLUVD | **(4)** | *0.5LVD | **(0.5)** | 0 | **(0)** |
| MM14 | 0 | **(0)** | 1RUVD 1.5LUVM | **(5)** | 1LUVD | **(2)** | 0 | **(0)** | 0 | **(0)** | 0 | **(0)** |
| MM15 | *0.5B | **(0.5)** | 1RVD 1LUVD | **(3)** | 1RVD 1LUVD | **(3)** | 1RLVD | **(2)** | 1RLVD | **(2)** | 1LVD | **(1)** |
| MM01 | 0 | **(0)** | 1LUVD | **(2)** | 0 | **(0)** | *1RCM | **(1)** | 0 | **(0)** | 0 | **(0)** |
| MM03 | 1RVM | **(1)** | 1RLVCD | **(4)** | 2.5LCVD 1RVD | **(6)** | 1RVD 1.5LCVD | **(4)** | 1RLVD | **(2)** | 1RLVD | **(2)** |
| MM05 | 1LCVD | **(2)** | 1LCVD | **(2)** | 0.5LVD | **(0.5)** | *1LUCVD | **(3)** | *1LUCVD | **(3)** | 0 | **(0)** |
| MM06 | 0.5LUCVD | **(1.5)** | 1LUCVD | **(3)** | 3LCVDM, 2RVM | **(8)** | 1LCVD | **(2)** | 1RVD 1LCVD | **(3)** | 1LVD | **(1)** |
| MM07 | 0 | **(0)** | 0 | **(0)** | 0 | **(0)** | *0.5RUD | **(0.5)** | *0.5RUD | **(0.5)** | 0 | **(0)** |
| MM08 | 1LVB | **(1)** | 1RVD 1LUVBD | **(3)** | 1LVB | **(1)** | 0 | **(0)** | 0 | **(0)** | 0 | **(0)** |

**Descriptive Score- Severity:** 0= normal, 1=mild bronchial or interstitial (mild increased soft tissue opacity), 2= moderate bronchial or interstitial (increased opacity with blurry vessels), 3=alveolar (unable to see vessels), **Location:** R=right, L=left, U=upper lung, V=lower lung, C=middle lung (right middle or caudal segment of left cranial lobe), **Other:** B=centered on airway/bronchi or hilar, P=peripheral, D=diffuse, M= multifocal or patchy, E=pleural effusion. **Total Score-** calculated by combining severity score with number of affected locations (possible range 0-18). [1]Baseline= day -7 to day 0 (challenge). [2]End of project=day 14-16 post infection. *Suspect opacity may be related to hypoinflation.

| Table S6. CT Scoring Infected Macaques, Pilot and Two Week Study | | | | | | | | | | | | |
|---|---|---|---|---|---|---|---|---|---|---|---|---|
| **Animal** | **Day 0** | | **Day 1** | | **Day 2** | | **Day 3** | | **Day 6** | | **Day 12-14** | |
| | Descriptive Score | **(Total)** | Descriptive Score | **(Total)** | Descriptive Score | **(Total)** | Descriptive Score | **(Total)** | Descriptive Score | **(Total)** | Descriptive Score | **(Total)** |
| **MM13** | 0 | **(0)** | 1RVN | **(1)** | 2RLVGN & 1RLUGD | **(6)** | 2RLUVG-BOPN | **(8)** | -- | - | -- | - |
| **MM16** | 0 | **(0)** | 3LVBN, 1RLUVGDP | **(7)** | 2RLUVDG | **(8)** | 1 RLVPG | **(2)** | -- | - | -- | - |
| **MM02** | 0.5RUVOP | **(1)** | 1RLUVOPG, RUVN | **(4)** | 1RUVOPG, RUVN | **(2)** | -- | - | -- | - | -- | - |
| **MM04** | 0 | **(0)** | 3LC,3RVO, 1RVB,1LVG | **(8)** | 3LC,3RVO, 2RVGB | **(8)** | -- | - | -- | - | -- | - |
| **MM03** | -- | - | -- | - | -- | - | -- | - | 3LVNPO, 1RVGD, 2LUCB | **(8)** | 2LVOP, 0.5RVG | **(2.5)** |
| **MM05** | -- | - | -- | - | -- | - | -- | - | 2RLUCVGD, RLUVA | **(12)** | 1RLUVGP, RLUVA,LVN | **(4)** |
| **MM06** | -- | - | -- | - | -- | - | -- | - | 1RLUCVDGP, 3RLCO, RLUCVA | **(12)** | 1RLUCVGP, 3RLCO, RLUCVA | **(12)** |
| **MM10** | -- | - | -- | - | -- | - | -- | - | 1RLUVPG, RVN | **(4)** | 0.5RVG, 0.5RVN | **(1)** |
| **MM11** | -- | - | -- | - | -- | - | -- | - | 1RLUVPG, RLVN | **(4)** | 0.5RLVG | **(1)** |
| **MM12** | -- | - | -- | - | -- | - | -- | - | 1.5RLUVPG, RUN | **(6)** | 1RLUVPG | **(4)** |

**Descriptive Score- Severity:** 0= normal, 1=mild bronchial or interstitial (ground glass opacity) 2= moderate bronchial or interstitial (dense ground glass opacity with crazy paving pattern), 3=alveolar (uniform soft tissue), **Location:** R=right, L=left, U=upper lung, V=lower lung, C=middle lung (right middle or caudal segment of left cranial lobe), **Other:** B=centered on airway/bronchi or hilar, O=peripheral, D=diffuse, P= multifocal or patchy, E=pleural effusion, G= ground glass, A=Bulla. **Total Score-** calculated by combining severity score with number of affected locations (possible range 0-18).

| Table S7. List of antibodies. | | | | | |
|---|---|---|---|---|---|
| Antibody | Supplier | Clone | Cat Number | Lot | Validation Statement: Reactivity |
| CD69 (FITC) | BD Bioscience | FN50 | 555530 | | Human (QC Testing) Rhesus, Cynomolgus, Baboon (Tested in Development) |
| CD4 (PCP-Cy5.5) | BD Bioscience | L200 | 552838 | | Rhesus, Cynomolgus, Baboon (QC Testing) Human (Tested in Development) |
| CD8 (APC) | Biolegend | RPA-T8 | 301049 | | Chimpanzee, Baboon, Cynomolgus, Rhesus, Pigtailed Macaque, Sooty Mangabey |
| CD3 (AL700) | BD Bioscience | SP34-2 | 557917 | | Rhesus, Cynomolgus, Baboon (QC Testing) Human (Tested in Development) |
| CD20 (APC-H7) | BD Bioscience | 2H7 | 560853 | | Rhesus, Cynomolgus, Baboon (QC Testing) Human (Tested in Development) |
| CD95 (BV 421) | BD Bioscience | DX2 | 562616 | | Human (QC Testing) Rhesus, Cynomolgus, Baboon (Tested in Development) |
| Ki67 (AF 488) | BD Bioscience | B56 | 558616 | | Human (QC Testing) Mouse (Tested in Development) Rat, Rhesus (Reported) |
| CCR7 (BV605) | BD Bioscience | 3D12 | 563711 | | Human (QC Testing) , Rhesus (NHP Reagent Resource) |
| CCR5 (PE) | BD Bioscience | 3A9 | 556042 | | Human (QC Testing) Rhesus, Cynomolgus (Tested in Development) |
| CD28 (PE-Cy7) | BD Bioscience | CD28.2 | 560684 | | Human (QC Testing), Rhesus (NHP Reagent Resource) |
| CD45 (BUV395) | BD Bioscience | D058-1283 | 564099 | | Rhesus, Cynomolgus, Baboon (QC Testing) |
| HLA-DR (APC-Cy7) | BD Bioscience | L243 | 335796 | | Human (QC Testing), Rhesus (NHP Reagent Resource) |
| PD-1 (BV421) | Biolegend | EH12.2H7 | 329920 | | Human, African Green, Baboon, Chimpanzee, Common Marmoset, Cynomolgus, Rhesus, Squirrel Monkey |
| CCR7 (BV480) | BD Bioscience | 3D12 | 566099 | | Human (QC Testing), Rhesus (NHP Reagent Resource) |
| CCR5 (BV650) | BD Bioscience | 3A9 | 564999 | | Human (QC Testing) Rhesus, Cynomolgus (Tested in Development) |
| CCR6 (BV711) | BD Bioscience | 11A9 | 563923 | | Human (QC Testing) Rhesus, Cynomolgus, Baboon (Tested in Development) |
| LAG-3 (PE) | R&D Systems | | FAB2319P | | Human, Rhesus (Validated in lab, Reported) |
| CXCR3 (PE-Cy7) | BD Bioscience | 1C6 | 560831 | | Human (QC Testing) Rhesus, Cynomolgus, Baboon (Tested in Development) |
| CD103 (FITC) | ThermoFisher | B-Ly7 | 11-1038-42 | | Human, Rhesus (NHP Reagent Resource) |
| CD123 (PerCP-Cy5.5) | BD Bioscience | 7G3 | 562391 | | Human (QC Testing) , Rhesus (NHP Reagent Resource) |
| CD14 (APC) | BD Bioscience | M5E2 | 561383 | | Rhesus, Cynomolgus, Baboon (QC Testing) |
| CD20 (AF700) | BD Bioscience | 2H7 | 560631 | | Human (QC Testing) Rhesus, Cynomolgus, Baboon (Tested in Development) |
| CD206 (BV421) | BD Bioscience | 19.2 | 564062 | | Human (QC Testing) , Rhesus (NHP Reagent Resource) |
| CD16 (BV480) | BD Bioscience | 3G8 | 566108 | | Human (QC Testing) Rhesus, Cynomolgus, Baboon (Tested in Development) |
| CD11c (BV711) | Biolegend | 3.9 | 301630 | | Human, African Green, Baboon, Chimpanzee, Cynomolgus, Rhesus, Squirrel Monkey |
| CD163 (PE) | BD Bioscience | GHI/61 | 556018 | | Human (QC Testing), Rhesus (NHP Reagent Resource) |
| CD40 (PE-Cy7) | BD Bioscience | 5C3 | 561215 | | Human (QC Testing) Rhesus, Cynomolgus, Baboon (Tested in Development) |
| IFN-Y (APC-Cy7) | Biolegend | B27 | **506524** | | Chimpanzee, Baboon, Cynomolgus, Rhesus, Pigtailed Macaque, African Green, Sooty Mangabey |
| IL-17 (BV605) | Biolegend | BL168 | 512326 | | Human, Rhesus (Validated in lab, Reported) |
| TNF-alpha (BV650) | Biolegend | MAb11 | 502938 | | Human, Cat (Feline)11 Cross-Reactivity: Chimpanzee, Baboon, Cynomolgus, Rhesus, Pigtailed Macaque, Sooty Mangabey, Swine (Pig, Porcine) |
| GrB (PE) | BD Bioscience | GB11 | 561142 | | **Human (QC Testing), Rhesus (NHP Reagent Resource)** |
| IL-2 (BUV737) | BD Bioscience | MQ1-17H12 | 612836 | | Human (QC Testing) Rhesus, Cynomolgus, Baboon (Tested in Development) |
